# Coordination of NDC80 and Ska complexes at the kinetochore-microtubule interface in human cells

**DOI:** 10.1101/820530

**Authors:** Robert Wimbish, Keith F. DeLuca, Jeanne E. Mick, Jack Himes, Ignacio J. Sánchez, A. Arockia Jeyaprakash, Jennifer G. DeLuca

## Abstract

The conserved kinetochore-associated NDC80 complex (comprised of Hec1/Ndc80, Nuf2, Spc24, and Spc25) has well-documented roles in mitosis including (1) connecting mitotic chromosomes to spindle microtubules to establish force-transducing kinetochore-microtubule attachments, and (2) regulating the binding strength between kinetochores and microtubules such that correct attachments are stabilized and erroneous attachments are released. Although the NDC80 complex plays a central role in forming and regulating attachments to microtubules, additional factors support these processes as well, including the spindle and kinetochore-associated (Ska) complex. Multiple lines of evidence suggest that Ska complexes strengthen attachments by increasing the ability of NDC80 complexes to bind microtubules, especially to depolymerizing microtubule plus-ends, but how this is accomplished remains unclear. Using cell-based and *in vitro* assays, we demonstrate that the Hec1 tail domain is dispensable for Ska complex recruitment to kinetochores and for generation of kinetochore-microtubule attachments in human cells. We further demonstrate that Hec1 tail phosphorylation regulates kinetochore-microtubule attachment stability independently of the Ska complex. Finally, we map the location of the Ska complex in cells to a region near the coiled-coil domain of the NDC80 complex, and demonstrate that this region is required for Ska complex recruitment to the NDC80 complex-microtubule interface.

## Introduction

Successful chromosome segregation during mitosis depends on the formation of stable attachments between chromosomes and spindle microtubules. These attachments are generated at kinetochores, which are macromolecular structures built on centromeric heterochromatin of mitotic chromosomes. Once stable kinetochore-microtubule connections are formed, forces generated by plus-end microtubule dynamics are harnessed for the purpose of congressing chromosomes to the spindle equator and silencing the spindle assembly checkpoint, which prevents anaphase onset until all kinetochores are properly attached to spindle microtubules. The kinetochore-associated NDC80 complex, comprised of the proteins Hec1 (also known as Ndc80), Nuf2, Spc24, and Spc25, serves as the core linkage between kinetochores and spindle microtubules (DeLuca and Musacchio, 2012). A direct interaction has been mapped between the “toe” domain of Hec1, which resides in its well-ordered, N-terminal calponin homology (CH) domain, and the microtubule lattice (Ciferri et al., 2008; Wilson-Kubalek et al., 2008; Alushin et al., 2010). This interaction is required for high affinity NDC80 complex-microtubule interactions *in vitro* and for kinetochore-microtubule attachment formation in cells from all organisms tested to date (Ciferri et al., 2008; Lampert et al., 2013; Sundin et al., 2011; Tooley et al., 2011; Cheerambathur et al., 2017). The Hec1 protein contains an N-terminal, unstructured “tail” domain that has also been implicated in forming kinetochore-microtubule attachments in cells, although requirement for the tail domain in this process varies among eukaryotic species. The Hec1 tail domain in *S. cerevisiae* and *C. elegans* is dispensable for formation of stable kinetochore-microtubule attachments (Kemmler et al., 2009; Demirel et al., 2012; Lampert et al., 2013; Cheerambathur et al., 2013). In contrast, expression of Hec1 mutants lacking the N-terminal tail domain in human or PtK1 cells impairs the formation of stable attachments (Guimaraes et al., 2008; Miller et al., 2008). The tail domain of Hec1 from all species tested, however, is required for high affinity binding of NDC80 complexes to microtubules *in vitro* (Wei et al., 2007; Ciferri et al., 2008; Miller et al., 2008; Lampert et al., 2013; Umbreit et al., 2012; Alushin et al., 2012; Cheerambathur et al., 2013; Zaytsev et al., 2015), suggesting that cellular factors likely compensate for Hec1 tail domain functions to varying degrees in different organisms.

In addition to generating attachments to spindle microtubules, kinetochores also regulate their stability. In early mitosis attachments are labile and undergo rapid turnover, whereas in late mitosis, attachments are stable and long-lived (Zhai et al., 1995; DeLuca et al., 2006; Cimini et al., 2006; Bakhoum et al. 2009). This scheme helps ensure that any erroneous attachments formed in early mitosis are released and corrected, and that mature attachments on correctly bi-oriented chromosomes are stabilized. The family of Aurora kinases is largely responsible for this phospho-regulation (Biggins et al., 1999; Tanaka et al., 2002; Carmena et al., 2012; Krenn and Musacchio, 2015), and the Hec1 N-terminal tail domain is an Aurora kinase target (DeLuca et al., 2006; Cheeseman et al., 2006). Nine sites in the tail domain have been identified as substrates of Aurora kinases A and B in vitro, and at least 5 are confirmed to be phosphorylated in cells (Nousiainen et al., 2006; DeLuca et al., 2011; Kettenbach et al., 2011; DeLuca et al., 2018). *In vitro*, progressive mutation of these 9 target sites to aspartic acid to mimic increasing phosphorylation results in a coordinate decrease in microtubule binding affinity of human NDC80 complexes (Zaytsev et al., 2015). Increasing the number of phospho-mimic substitutions also results in a corresponding decrease in kinetochore-microtubule attachment stability in mammalian cells (Zaytsev et al., 2014). Conversely, expression of Hec1 mutants in which all mapped Aurora kinase target sites are mutated to alanine to prevent phosphorylation results in hyper-stabilization of kinetochore-microtubule attachments and defective attachment error correction in mammalian cells (DeLuca et al., 2011; Sundin et al., 2011; Zaytsev et al., 2014; Tauchman et al., 2015; Long et al., 2017; Yoo et al., 2018). A similar phenomenon is observed in embryonic *C. elegans* cells, where mutation of the four mapped Hec1 tail domain Aurora kinase target sites to alanine results in premature kinetochore-microtubule stabilization (Cheerambathur et al., 2017). One model to explain these results proposes that increased Aurora kinase phosphorylation of the Hec1 tail decreases the affinity of the NDC80 complex for microtubules, which in turn decreases kinetochore-microtubule stability.

In addition to the NDC80 complex, the spindle and kinetochore associated (Ska) complex, a trimer comprised of Ska1, Ska2, and Ska3, contributes to the generation and stabilization of kinetochore-microtubule attachments. The Ska complex loads progressively onto kinetochores during mitosis and is required for efficient chromosome congression and for silencing the spindle assembly checkpoint (Hanisch et al., 2006; Gaitanos et al., 2009; Theis et al., 2009; Raaijmakers et al., 2009; Daum et al., 2009; Guimaraes and DeLuca, 2009; Sivakumar et al., 2014; Sivakumar et al., 2016; Auckland et al., 2017). The Ska complex binds both the NDC80 complex and microtubules, and stabilizes NDC80 complex-mediated kinetochore-microtubule attachments, likely through its ability to remain bound to depolymerizing microtubule plus-ends (Welburn et al., 2009; Jeyaprakash et al., 2012; Schmidt et al., 2013; Abad et al., 2014; Zhang et al., 2017; Helgeson et al., 2018). A major outstanding question is how the Ska complex is recruited to kinetochore-bound NDC80 complexes to promote kinetochore-microtubule attachment stability. Previous studies have suggested that this recruitment is mediated through the Hec1 tail domain (Cheerambathur et al., 2017; Janczyk et al., 2017), the Hec1 loop domain (Zhang et al., 2012; Zhang et al., 2017), and the coiled-coil regions of the hetero-tetrameric complex (Helgeson et al., 2018), thus the recruitment mechanism remains unresolved.

The Ska complex has also been implicated in regulating kinetochore-microtubule attachment stability. Expression of a non-phosphorylatable Hec1 tail domain mutant in *C. elegans* embryos resulted in increased recruitment of the Ska complex, whereas expression of a phospho-mimetic Hec1 mutant led to the opposite effect (Cheerambathur et al., 2017). Furthermore, increased stability of kinetochore-microtubule attachments observed in cells expressing the non-phosphorylatable mutant version of Hec1 was dependent on the presence of the Ska complex. Thus in some organisms, rather than directly regulating NDC80 complex-microtubule affinity, phosphorylation of the Hec1 tail likely controls the recruitment of Ska complexes, which in turn regulates attachment stability. Whether this mechanism functions in human cells remains to be tested.

Here we investigate how the human Ska complex is recruited to the NDC80 complex in cells and *in vitro*, and how Hec1 tail phosphorylation impacts Ska function. We report that the N-terminal Hec1 tail domain, while required for proper chromosome congression, is not explicitly required for either kinetochore-microtubule attachment or Ska complex recruitment to kinetochores in human cells. The tail domain is also dispensable for Ska complex-mediated enhancement of NDC80 complex-microtubule binding *in vitro*. We demonstrate that phospho-regulation of kinetochore-microtubule attachments occurs in the absence of the Ska complex in human cells, providing support for a mechanism whereby Aurora kinase phosphorylation of the Hec1 tail directly modulates kinetochore-microtubule attachment strength. Finally, using two color fluorescence localization microscopy, we map the location of the Ska complex to a region coincident with the central coiled-coil domains of the NDC80 complex and consistent with this, we find that microtubule binding of NDC80 complexes lacking the central coiled-coil domains is not enhanced by addition of Ska complexes.

## Results

### Phosphorylation of the Hec1 tail affects Ska complex loading to kinetochores

To determine how phosphorylation of the Hec1 tail impacts recruitment of Ska complexes to kinetochores, we expressed mutant versions of Hec1 in human cells in which the 9 mapped Aurora phosphorylation sites were mutated to either alanine (9A) to prevent phosphorylation, or to aspartic acid (9D) to mimic constitutive phosphorylation. We confirmed that expression of the exogenous constructs led to depletion of endogenous Hec1 protein from kinetochores to undetectable levels by staining cells with an antibody to phosphorylated Hec1 Serine 69 (pS69), which does not recognize 9A-or 9D-Hec1 proteins and whose levels do not vary during mitotic progression (DeLuca et al., 2018) (Fig. 1 A and B). Similar to the situation described for *C. elegans* (Cheerambathur et al., 2017), we found that kinetochores in cells expressing 9A-Hec1-GFP were enriched for the Ska complex, while kinetochores in cells expressing 9D-Hec1-GFP exhibited lower levels compared to kinetochores in cells expressing WT-Hec1-GFP (Fig. 1 A and C). Similar results were observed in cells expressing Hec1-GFP constructs and depleted of endogenous Hec1 by siRNA (Fig. S1 A-C). Although these results suggest that the phosphorylation state of the tail domain might directly regulate Ska complex recruitment to kinetochores, there is an important caveat to this experiment. Cells expressing 9A-Hec1 mutants generate hyper-stable kinetochore-microtubule attachments, in which kinetochore-microtubule bundle densities are increased (Zaytsev et al., 2014), the pulling forces between two sister kinetochores are higher (DeLuca et al., 2011; Yoo et al., 2018), and end-on kinetochore-microtubule attachments are formed earlier than in control cells (Fig. 1 D and E). Conversely, cells expressing 9D-Hec1 mutants fail to form stable kinetochore-microtubule attachments during mitosis (DeLuca et al., 2011; Zaytsev et al., 2014). Since the Ska complex loads to kinetochores as microtubule attachments are progressively stabilized (Hanisch et al., 2006; Auckland et al., 2017) (Fig. 1 F and G), results from the experiment described above (Fig. 1 A-C) do not allow us to differentiate between the two following scenarios: (1) dephosphorylation of the Hec1 tail promotes Ska complex recruitment, and in turn, the Ska complex increases kinetochore-microtubule attachment stability, or (2) dephosphorylation of the Hec1 tail generates stable kinetochore-microtubule attachments, and in turn, stable attachments promote recruitment of the Ska complex to kinetochores.

**Figure 1.**
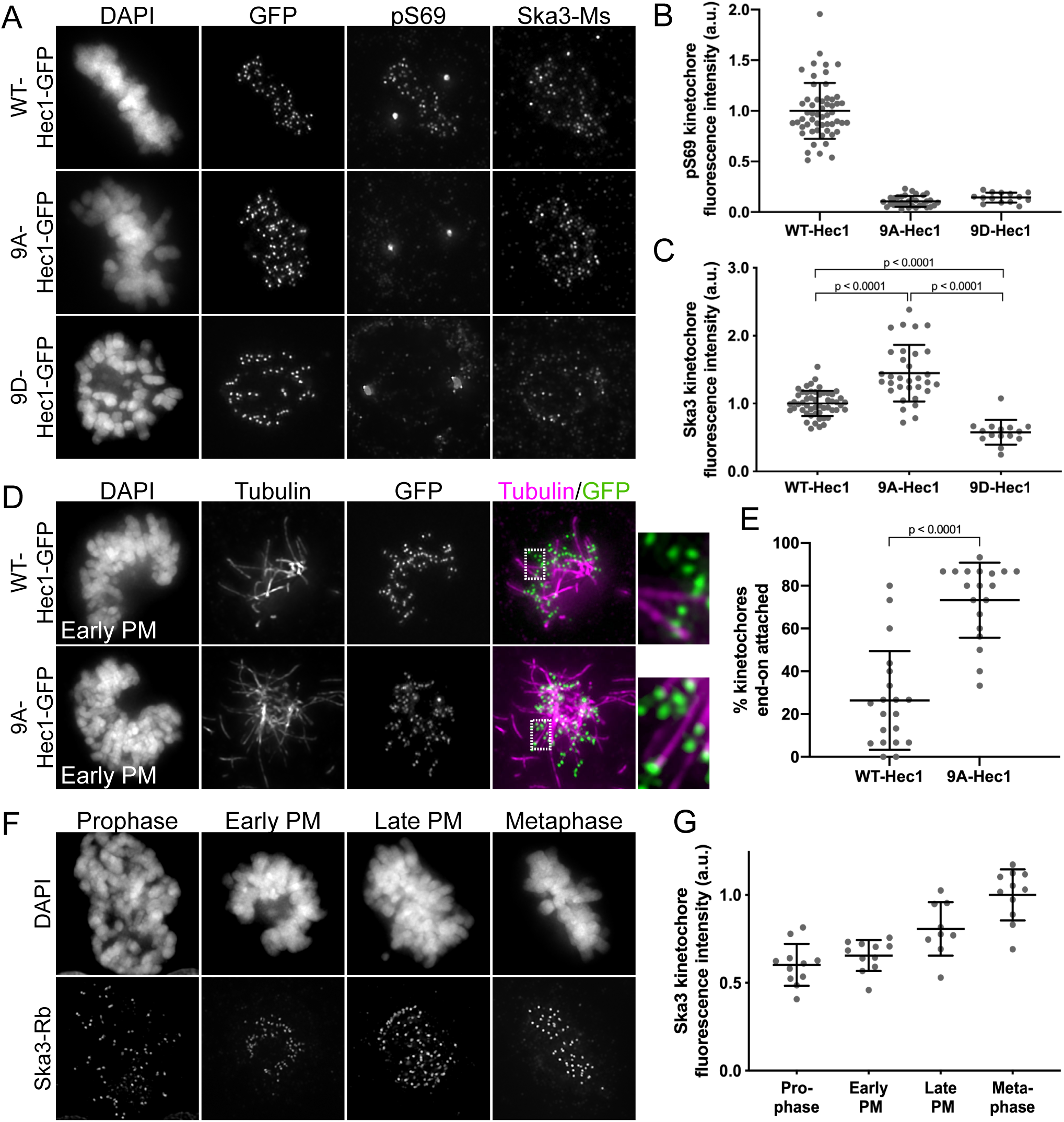
Phosphorylation of the Hec1 tail domain affects kinetochore-microtubule attachment stability and Ska complex loading to kinetochores. (A) Immunofluorescence images of cells expressing WT-, 9A-, and 9D-Hec1-GFP. Cells were fixed and stained using antibodies to Hec1 pS69 and Ska3 (generated in mouse). (B) Quantification of pS69 kinetochore fluorescence intensity from cells expressing WT, 9A-, and 9D-Hec1-GFP. For each condition, at least 20 kinetochores per cell were measured from at least 5 cells per experiment from 3 separate experiments. (C) Quantification of Ska3 kinetochore fluorescence intensity from cells expressing WT-, 9A-, and 9D-Hec1-GFP. For each condition, at least 20 kinetochores per cell were measured from at least 5 cells per experiment from 3 separate experiments. Statistical significance was determined by a one-way Anova analysis. (D) Immunofluorescence images of cold-treated cells expressing WT-and 9A-Hec1-GFP. Cells were incubated in cold DMEM on ice for 12 minutes prior to fixation, permeabilized, fixed and stained using antibodies to tubulin. Insets are enlargements of the region indicated by the dashed box. (E) Quantification of end-on attachment in cold-treated cells expressing WT-and 9A-Hec1-GFP. For each condition, at least 15 kinetochores per cell were measured from at least 9 cells per experiment from two separate experiments. A Student’s t-test was carried out to determine statistical significance. (F) Immunofluorescence images of untreated, control cells in different stages of mitosis fixed and stained with antibodies to Ska3 (generated in rabbit). (G) Quantification of Ska3 kinetochore fluorescence intensity in control cells in progressive stages of mitosis. For each mitotic phase, at least 20 kinetochores were measured from at least 4 cells per experiment from 2 separate experiments. For all graphs, each point on the scatter plot indicates the average value of all measured kinetochores from an individual cell.

### Phosphorylation of the Hec1 tail does not affect microtubule-independent Ska complex loading to kinetochores

To begin to differentiate between the two possibilities, we measured Ska complex loading to kinetochores in cells expressing WT, 9D-, and 9A-Hec1-GFP in the absence of microtubules. This allowed us to test how mutations in Hec1 affect Ska recruitment without the confounding effects of their impact on kinetochore-microtubule attachment stability. Previous reports have demonstrated that while Ska complexes are maximally loaded onto kinetochores after microtubule attachment, there is a population of microtubule-independent, Hec1-dependent Ska complex recruitment to kinetochores (Chan et al., 2012; Zhang et al., 2017). Cells transfected with either WT-, 9A-, or 9D-Hec1-GFP were synchronized and arrested in G2 with RO-3306 and then washed out into nocodazole prior to entry into mitosis. We confirmed that microtubule-independent Ska complex recruitment to kinetochores required the NDC80 complex (Fig. 2 A and B), and found that kinetochores in cells expressing WT-, 9D- or 9A-Hec1-GFP all loaded similar levels of the Ska complex (Fig. 2 C and D). These results suggest that in the absence of microtubules, the phosphorylation state of the human Hec1 tail domain does not influence Ska complex recruitment to kinetochores. In this experiment, all cells subjected to analysis entered mitosis in the presence of nocodazole, and therefore kinetochores had no contact with microtubules prior to fixation.

**Figure 2.**
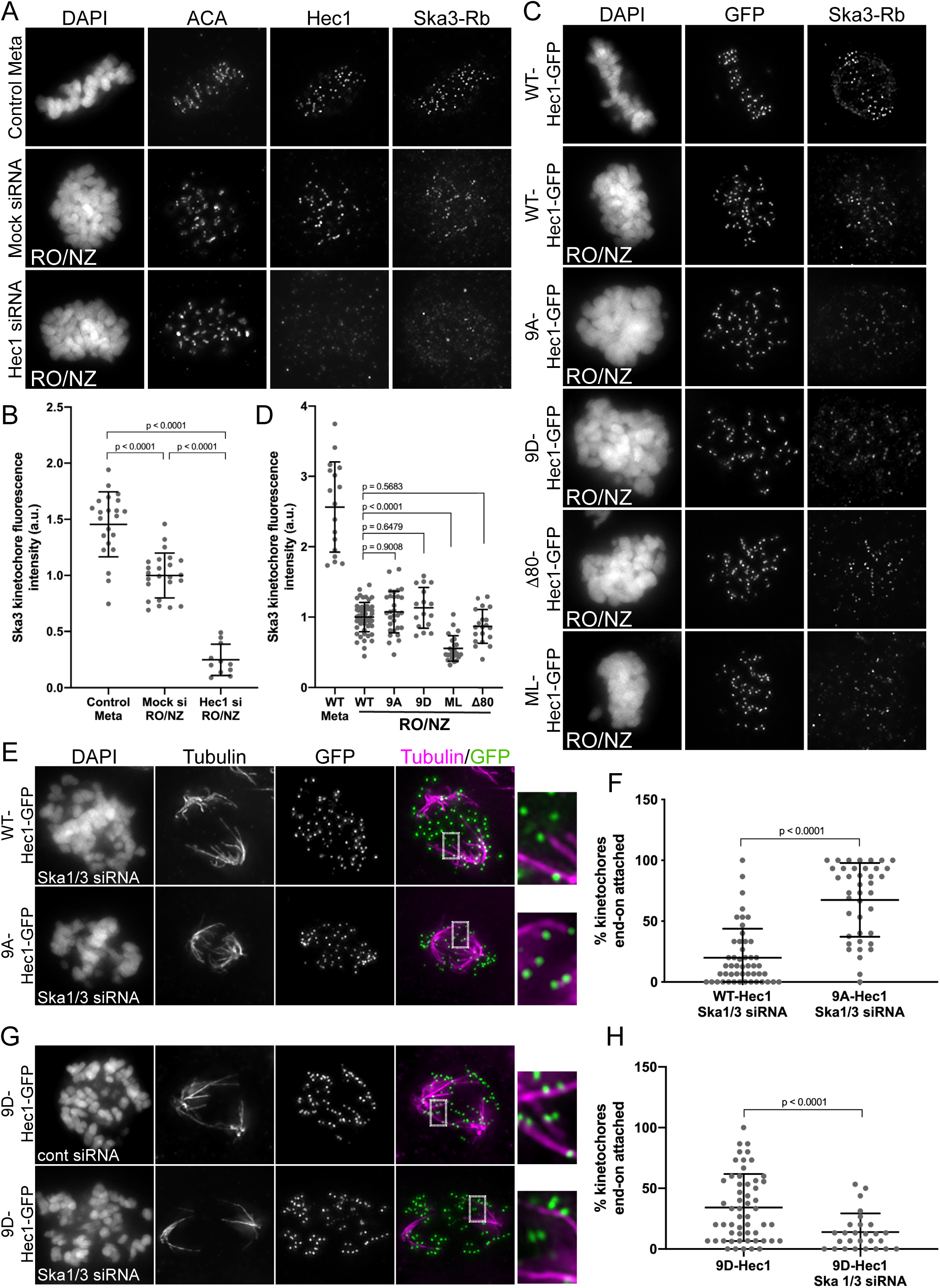
Hec1 tail dephosphorylation does not affect microtubule-independent Ska complex kinetochore loading and stabilizes attachments in the absence of the Ska complex. (A) Immunofluorescence images of untreated cells (top row) or RO3306-synchronized cells released into mitosis in the presence of 10 uM nocodazole (bottom two rows). Cells were stained with ACA (anti-centromere) antibodies and antibodies to Hec1 and Ska3 (rabbit). (B) Quantification of Ska3 kinetochore fluorescence intensity from cells described in panel (A). For each condition, at least 20 kinetochores per cell were measured from at least 5 cells per experiment from two separate experiments. Ska3 intensity was only measured for Hec1 siRNA-treated cells with <20% of endogenous kinetochore-associated Hec1 as determined by staining with an antibody to the CH domain of Hec1 (9G3). A Student’s t-test was carried out to determine statistical significance. (C) Immunofluorescence images of cells expressing the indicated Hec1-GFP fusion protein in the absence (top row) or presence of RO3306-synchronization and release into 10 µm nocodazole (remaining rows). Cells were stained with antibodies to Ska3 (rabbit). (D) Quantification of Ska3 kinetochore fluorescence intensity from cold-treated cells described in panel (C). For each condition, at least 20 kinetochores per cell were measured from at least 5 cells per experiment from 3 separate experiments. Statistical significance was determined by a one-way Anova analysis between RO3306-synchronized WT-Hec1-GFP expressing cells and the indicated Hec1 fusion proteins. (E) Immunofluorescence images of cold-treated cells expressing WT- and 9A-Hec1-GFP and treated with Ska1 and Ska3 siRNA. Cells were incubated in cold DMEM on ice for 12 minutes prior to fixation, permeabilized, fixed, and stained using antibodies to tubulin. Insets are enlargements of the region indicated by the dashed box. (F) Quantification of end-on attachment in cells expressing WT- and 9A-Hec1-GFP and treated with Ska1 and Ska3 siRNA. For each condition, at least 15 kinetochores were measured from at least 10 cells from 3 separate experiments. A Student’s t-test was carried out to determine statistical significance. (G) Immunofluorescence images of cells expressing 9D-Hec1-GFP and treated with (bottom panel) or without (top panel) Ska1 and Ska3 siRNA. Cells were incubated in cold DMEM on ice for 12 minutes, permeabilized, fixed, and stained using antibodies to tubulin. Insets are enlargements of the region indicated by the dashed box. (H) Quantification of end-on attachments in cold-treated cells expressing 9D-Hec1-GFP and treated with or without Ska1 and Ska3 siRNA. For each condition, at least 15 kinetochores were measured per cell from at least 9 cells per experiment from at least 3 separate experiments. A Student’s t-test was carried out to determine statistical significance.

Interestingly, when we carried out a similar experiment in an asynchronous population, where nocodazole was added to cells in various stages of mitosis, we found that kinetochores in cells expressing 9A-Hec1-GFP exhibited higher levels of Ska3 compared to those in cells expressing WT-Hec1-GFP (Fig. S2 A and B). We speculate that a population of kinetochores in asynchronous cells expressing 9A-Hec1-GFP had previously established kinetochore-microtubule attachments and loaded high levels of the Ska complex to kinetochores prior to exposure to nocodazole. These results suggest that once Ska complexes are loaded onto kinetochores by microtubule attachment, a sub-population of the complex remains bound even after microtubule depolymerization.

### Hec1 tail phosphorylation contributes to kinetochore-microtubule attachment stability independently of the Ska complex

To investigate the functional dependencies between Hec1 tail dephosphorylation, Ska complex loading, and kinetochore-microtubule attachment stability in human cells, we tested if stable attachments formed in human cells expressing 9A-Hec1-GFP were dependent on the Ska complex. Cells were depleted of Ska1 and Ska3, transfected with WT- or 9A-Hec1-GFP constructs, subjected to cold-treatment, fixed and stained with tubulin antibodies, and scored for the presence of end-on kinetochore-microtubule attachments. As expected, cells depleted of Ska1 and Ska3 exhibited defects in chromosome alignment and end-on kinetochore-microtubule attachment formation (Fig. S2 C and D). Similarly, in cells depleted of Ska1/Ska3 and expressing WT-Hec1-GFP, we observed defects in chromosome alignment and formation of end-on kinetochore-microtubule attachments (Fig. 2 E and F). However, in Ska1/Ska3-depleted cells expressing 9A-Hec1-GFP, we observed robust formation of end-on attachments similar to what we observed in non-Ska1/3 depleted cells (Fig. 2 E and F), suggesting that Hec1 tail phosphorylation and Ska complex recruitment contribute to regulation of kinetochore-microtubule attachments independently of each other. We reasoned that if this were the case, then the destabilizing effects of expressing a phospho-mimetic Hec1 tail mutant in Ska 1/3-depleted cells should be more severe than the effects of expressing a phospho-mimetic Hec1 tail mutant in non-Ska1/3-depleted cells. We found that cells expressing 9D-Hec1-GFP exhibited defects in forming stable, end-on kinetochore-microtubule attachments (average of ∼34% of kinetochores attached per cell), which is consistent with previous studies (Fig. 2 G and H). In cells depleted of Ska1 and Ska3, expression of 9D-Hec1-GFP indeed resulted in a more severe kinetochore-microtubule attachment defect (average of ∼14% of kinetochores attached per cell) (Fig. 2 G and H), providing further evidence that in human cells, phosphorylation of the Hec1 tail contributes to kinetochore-microtubule attachment stability independently of the Ska complex.

### The Hec1 tail domain is not required for Ska complex-mediated enhancement of NDC80 complex-microtubule binding

Independent of its phosphorylation state, the tail domain of Hec1 has been implicated in recruiting the Ska complex to the NDC80 complex-microtubule interface and to kinetochores in human cells (Janczyk et al., 2017). To further investigate this, we first asked if the tail domain is required *in vitro* for Ska complexes to enhance NDC80 complex-microtubule affinity. Previous studies have shown that purified, recombinant Ska complexes increase the affinity of NDC80 complexes for microtubules *in vitro* (Schmidt et al., 2012; Helgeson et al., 2018). We therefore measured the microtubule binding affinity of GFP-tagged, recombinantly-expressed, purified NDC80 complexes containing WT-Hec1 and Hec1 deleted of its N-terminal 80 amino acid tail domain (Δ80-Hec1) using a TIRF-based fluorescence assay. For these experiments, we generated NDC80 complexes in which Nuf2 is fused to Spc24 and Hec1 is fused to Spc25-GFP (Fig. 3 A), termed NDC80^Bronsai^. These complexes are missing the tetramerization domains from all four subunits, but contain the majority of the central coiled-coil region of the complex (Ciferri et al., 2005; Ciferri et al., 2008), as well as the “loop” domain of Hec1, which is a 40 amino acid region that briefly disrupts the coiled-coil region (Maiolica et al., 2007). The name represents a hybrid between “NDC80^Bonsai^,” which is an engineered, truncated version of the NDC80 complex comprised of a Nuf2-Spc24 fusion and an Ndc80/Hec1-Spc25-GFP fusion missing the central coiled-coil and tetramerization domains (Ciferri et al., 2008) and “NDC80^Broccoli^,” which is a dimer of nearly full-length Nuf2 and Ndc80/Hec1 containing the coiled-coil and loop domains (Schmidt et al., 2012). For the binding assays, we incubated increasing concentrations of GFP-labeled NDC80^Bronsai^ complexes with Alexa647-labeled microtubules in the presence or absence of 10 nM recombinantly expressed human Ska complex, and measured the average fluorescence intensity along microtubules. WT-NDC80^Bronsai^ complexes robustly bound microtubules and binding was increased upon the addition of Ska complexes (Fig. 3 A and B). For WT-NDC80^Bronsai^, we measured approximately a 2-fold increase in binding affinity after Ska complex addition (Fig. 3 B). The Δ80-NDC80^Bronsai^ complexes bound to microtubules with significantly lower affinity than the WT complexes (Fig. 3 C and D), which is consistent with previously published studies (Miller et al., 2008; Umbreit et al., 2012; Zaytsev et al., 2015). However, addition of purified Ska complex significantly increased the affinity of the poorly-binding Δ80-NDC80^Bronsai^ complexes for microtubules (Fig. 3 D). These results demonstrate that the Hec1 tail domain is not required for Ska complex-mediated enhancement of microtubule binding by NDC80 complexes *in vitro*, and that Ska complexes are able to compensate for the weak microtubule binding observed with NDC80 complexes lacking the N-terminal tail domain. These results also demonstrate that the tetramerization domain of the NDC80 complex is not required for Ska complex binding. We note that one difference between the TIRF-based microtubule binding assays described here and those described in our previous study (Zaytsev et al., 2015), is the choice of assay buffer. When we used standard microtubule binding assay buffers BRB80 (80 mM PIPES, 1 mM MgCl_2_, 1 mM EGTA, pH 6.8) or BRB20 (20 mM PIPES, 1 mM MgCl_2_, 1 mM EGTA, pH 6.8), purified Ska complexes aggregated in the presence of microtubules (Fig. S3 A). In addition, Ska complexes induced aggregation of NDC80 complexes on microtubules in the presence of BRB80 (Fig. S3 B), which precluded quantitative analysis of fluorescence intensities along microtubules. We therefore developed “SN” buffer (for “Ska-NDC80”) for our assays (20 mM TRIS, 50 mM NaCl, 6 mg/ml BSA, 4 mM DTT, pH 7.0), which did not induce aggregation of either Ska or NDC80 complexes (Fig. S3 A).

**Figure 3.**
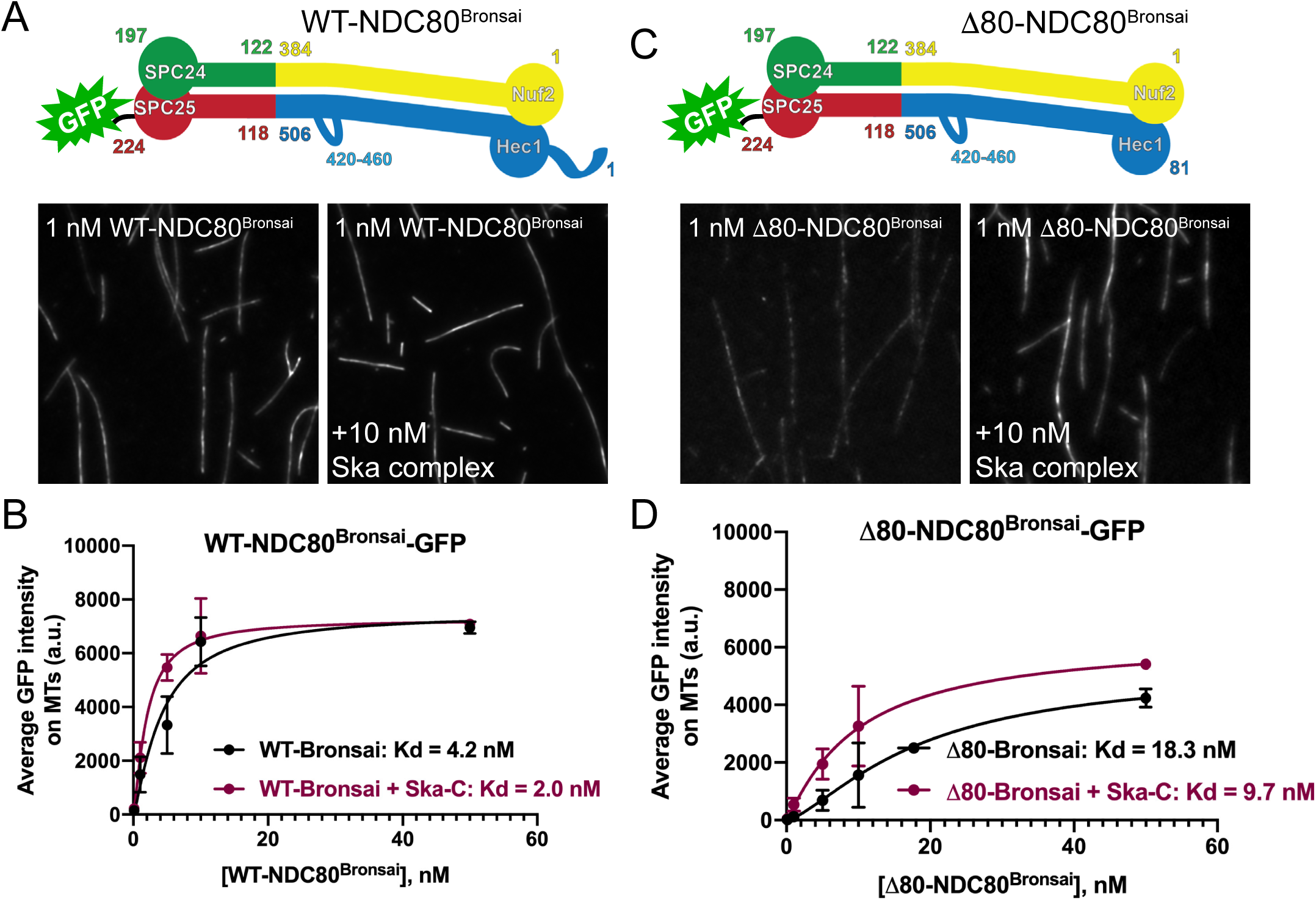
The Hec1 tail domain is not required for Ska complex-mediated enhancement of NDC80 complex-microtubule binding. (A and C) Top: Schematics of NDC80^Bronsai^ complexes used in the TIRF-based microtubule binding experiments. Bottom: GFP fluorescence images of NDC80 complexes decorating microtubules in the presence and absence of Ska complex. All images show a single concentration of the NDC80 complex from the experiment (1 nM) with and without added Ska complex (10 nM). (B and D) Binding curves from the microtubule binding assays. Datapoints and curve fits shown in black are from experiments without added Ska complex. Those shown in burgundy are from experiments with added Ska complex. Each point on the curve represents the average fluorescence intensity from three separate experiments. At each concentration, GFP-NDC80 complex fluorescence intensity was measured from at least 40 individual microtubules from at least 10 different TIRF fields per experiment.

### The Hec1 tail domain is not required for Ska complex recruitment to kinetochores or for kinetochore-MT attachment in human cells

In light of our *in vitro* results, we next wanted to ask if the Hec1 tail domain is required for Ska complex recruitment to kinetochores in human cells. For these experiments, we expressed exogenous Δ80-Hec1-GFP in HeLa cells and analyzed only cells with undetectable levels of endogenous Hec1 at kinetochores. Previous studies in mammalian cells demonstrated that Hec1 tail deletion impacts kinetochore-microtubule attachment stability as evidenced by reduction in inter-kinetochore distances, decreased cold-resistant microtubule attachments, failure to align chromosomes, and significant mitotic delays (Guimaraes et al., 2008; Miller et al., 2008; Etemad et al., 2015; Janczyk et al., 2017). In line with this, we found that cells expressing Δ80-Hec1-GFP exhibited significant chromosome alignment defects and decreased inter-kinetochore distances (Fig. 4 A and B, Fig S4 A and B). However, contrary to previous studies, we found that cells expressing Δ80-Hec1-GFP were competent to form cold-resistant kinetochore-microtubule attachments (Figure 4 C). This is in contrast to cells expressing 9D-Hec1, which are neither able to properly align chromosomes or form stable, cold-resistant kinetochore-microtubule attachments (Fig. 4 A-C; DeLuca et al., 2011). Analysis of spindle morphology in Δ80-Hec1-expressing cells revealed that the majority of cells with unaligned chromosomes contained multi-polar spindles (Figure 4 D). To further investigate this phenotype, we carried out time-lapse imaging of mCherry-tubulin and Hec1-GFP expressing cells. In the case of WT-Hec1-GFP expressing cells, we found that almost all cells formed bi-polar spindles and entered anaphase without errors, with only ∼10% of cells undergoing spindle fragmentation prior to anaphase. Strikingly, while the majority of Δ80-Hec1-GFP expressing cells initially formed bi-polar spindles, ∼50% of cells eventually exhibited spindle pole fragmentation and loss of chromosome alignment (Figure 4 E and F). This phenotype was similar to cells expressing 9A-Hec1-GFP, which also exhibited spindle pole fragmentation during mitotic progression. We hypothesized that cells expressing Δ80-Hec1-GFP may have lost their ability to regulate kinetochore-microtubule attachment formation through Aurora kinase phosphorylation, which might contribute to the observed phenotypes. To test this hypothesis, we analyzed cold-resistant end-on attachments in early prometaphase cells shortly after nuclear envelope breakdown. Similar to 9A-Hec1-GFP expressing cells, and in contrast to WT-Hec1-GFP expressing cells, early prometaphase cells expressing Δ80-Hec1-GFP formed robust end-on kinetochore-microtubule attachments that resisted cold depolymerization (Figure 4 G and H), suggesting that attachments were formed prematurely in these cells, likely due to loss of negative regulation by Aurora kinases. However, in contrast to cells expressing 9A-Hec1-GFP, cells expressing Δ80-Hec1-GFP were unable to generate wild-type levels of tension across bi-oriented sister kinetochore pairs in later mitosis, as evidenced by lower inter-kinetochore distances (Figure S4 A and B). This result suggests that while the Hec1 tail is not explicitly required for kinetochore-microtubule attachment formation, it may be an important factor for generating force at the attachment interface.

**Figure 4.**
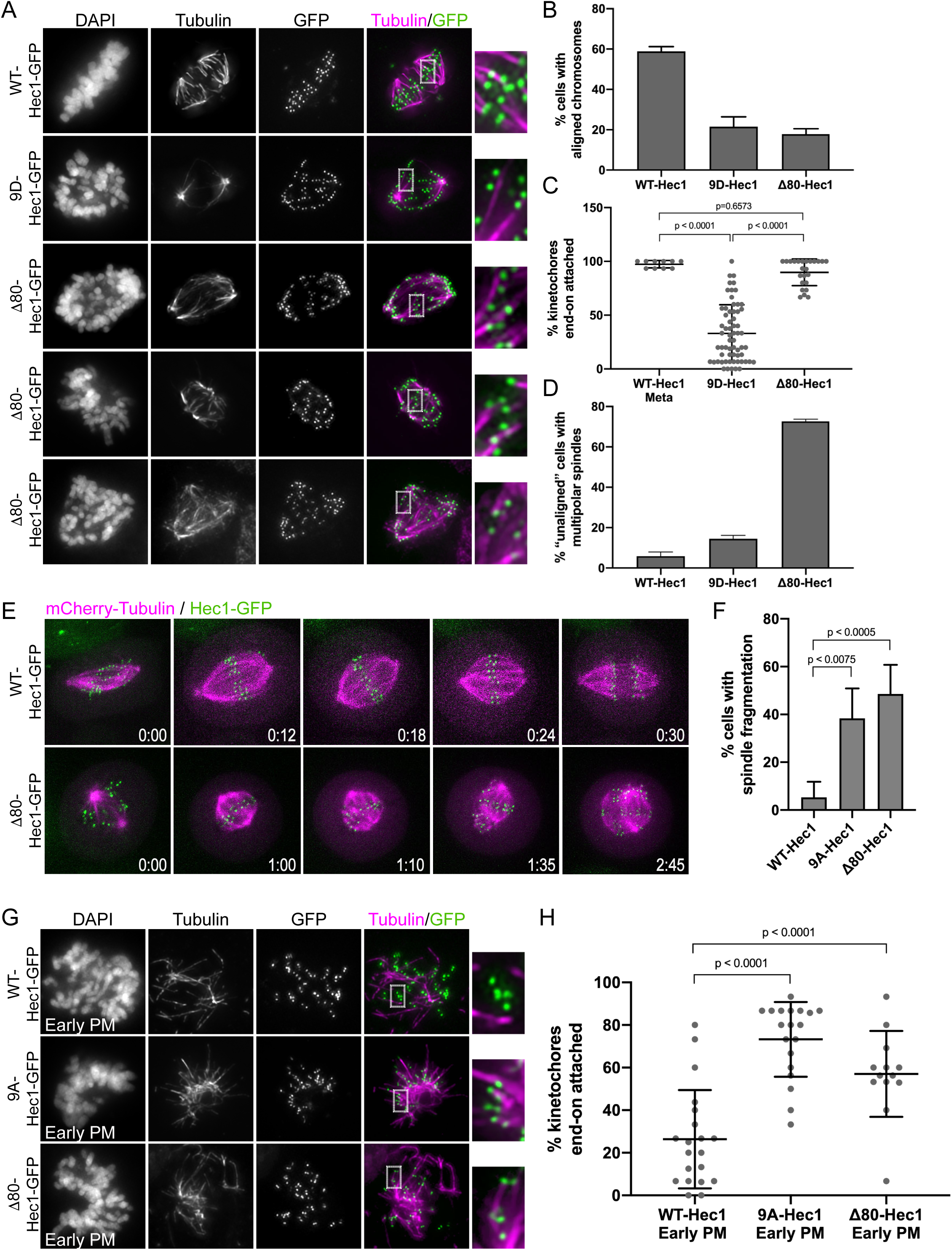
The Hec1 tail domain is not required for the formation of stable end-on kinetochore-microtubule attachments in cells. (A) Immunofluorescence images of cold-treated cells expressing WT-, 9D-, and Δ80-Hec1-GFP. Cells were incubated in cold DMEM for 12 minutes prior to fixation, permeabilized, fixed and stained using antibodies to tubulin. Insets are enlargements of the region indicated by the dashed box. Three examples of cells expressing Δ80-Hec1-GFP are shown. (B) Quantification of chromosome alignment in cells expressing WT-, 9D-, and Δ80-Hec1-GFP. For each condition, chromosome alignment was assessed in at least 100 cells per experiment from 2 separate experiments. Cells were scored as “aligned” if they had a metaphase plate with <5 chromosomes off the plate. (C) Quantification of end-on attachment in cells expressing WT-, 9D-, and Δ80-Hec1-GFP and cold-treated prior to fixation. For each condition, at least 15 kinetochores per cell were measured from at least 10 cells per experiment from 2 separate experiments. Statistical significance was determined by a one-way Anova analysis. (D) Quantification of multipolarity observed in cells expressing WT-, 9D- and Δ80-Hec1-GFP. Cells with unaligned chromosomes were scored for containing bi-vs multi-polar spindles, and the percent of cells with multipolar spindles is shown. For each condition, at least 100 cells per experiment were analyzed from 2 separate experiments. (E) Still images from time-lapse experiments of cells expressing Hec1-GFP and mCherry-tubulin. Cells expressing Δ80-Hec1-GFP (bottom panel) exhibit spindle pole fragmentation and loss of chromosome alignment. (F) Quantification of spindle pole fragmentation frequency quantified from time-lapse imaging experiments. Cells were scored as undergoing fragmentation events if loss of spindle bipolarity was observed during time-lapse imaging as determined from mCherry-tubulin signal. Quantifications shown are averages from at least three independent experiments. Total cell numbers assessed is as follows: 46, 27, and 47, for cells expressing WT-, 9A, and Δ80-Hec1-GFP, respectively. Statistical significance was determined by a one-way Anova analysis. (G) Immunofluorescence images of cold-treated, early prometaphase cells expressing WT, 9A-, and Δ80-Hec1-GFP. Cells were incubated in cold DMEM on ice for 12 minutes, permeabilized, fixed, and stained with antibodies to tubulin. Insets are enlargements of the region indicated by the dashed box. (H) Quantification of end-on attachment in early prometaphase cells expressing WT-, 9A-, and Δ80-Hec1-GFP. The WT- and 9A-Hec1 data shown are from the experiment presented in Figure 2. For each condition, at least 15 kinetochores per cell were measured from at least 6 cells per experiment from at least 2 separate experiments. Statistical significance was determined by a one-way Anova analysis.

We were somewhat surprised at the ability of cells expressing Δ80-Hec1-GFP to retain end-on attachments after cold-treatment, since it has been previously observed that the tail domain contributes to the formation and/or maintenance of kinetochore-microtubule attachments in both human and marsupial cells (Miller et al., 2008; Guimaraes et al., 2008; Etemad et al., 2015; Janczyk et al., 2017). To confirm that this was not a cell-type specific phenomenon, we expressed WT- and Δ80-Hec1-GFP constructs in human RPE1 cells and found that, similar to what was observed in HeLa cells, RPE1 cells expressing Δ80-Hec1-GFP were competent to form cold-resistant end-on kinetochore-microtubule attachments (Fig. S5 D). Interestingly, we found that location of the GFP had a major impact on the ability of Δ80-Hec1-expressing cells to form kinetochore-microtubule attachments. HeLa cells expressing either C- or N-terminally GFP-tagged WT-Hec1 constructs formed stable, end-on attachments, as previously reported (Miller et al., 2008; Guimaraes et al., 2008; DeLuca et al., 2011; Etemad et al., 2015; Janczyk et al., 2017). In contrast, HeLa cells expressing C-terminally GFP-tagged Δ80-Hec1 formed end-on kinetochore-microtubule attachments, while those expressing the N-terminally GFP-tagged Δ80-Hec1 did not (Fig. S5 A). Similar results were found in RPE1 cells (Fig. S5 D). We quantified attachment stability in HeLa cells expressing either C- or N-terminally GFP-tagged Δ80-Hec1 using a cold-induced microtubule depolymerization assay, and confirmed that while cells expressing C-terminally tagged Δ80-Hec1 were able to form cold-stable, end-on attachments, cells expressing N-terminally tagged Δ80-Hec1 were not (Figure S5 B and C).

After characterizing the phenotype of cells expressing Δ80-Hec1-GFP, we returned to our original question and measured Ska3 levels at kinetochores in cells expressing tail-less Hec1. These experiments revealed no significant difference in Ska3 levels at kinetochores in cells expressing WT-vs. Δ80-Hec1-GFP (Fig. 5 A and B). Microtubule-independent Ska3 recruitment to kinetochores also remained high in cells expressing Δ80-Hec1-GFP (Fig. 2 C and D), suggesting that the tail domain is dispensable for both microtubule-dependent and -independent Ska complex recruitment. Given that purified Ska complexes compensated for the weak binding affinity of Δ80-NDC80 complexes in vitro (Fig. 3), we asked if formation of kinetochore-microtubule attachments in cells expressing Δ80-Hec1-GFP required the presence of an in-tact Ska complex. We depleted Ska complex components Ska1 and Ska3 from HeLa cells, expressed either WT- or Δ80-Hec1-GFP, incubated cells in cold media, and measured the abundance of end-on kinetochore-microtubule attachments. Kinetochore-microtubule attachments failed to form in Ska1/3-depleted cells expressing either WT- or Δ80-Hec1-GFP (Fig. 5 C and D), which is in contrast to cells expressing 9A-Hec1-GFP (Fig. 2 E and F). Thus, tail-less NDC80 complexes, similar to WT complexes, require the Ska complex to form attachments to microtubules. Collectively, our results suggest that the tail domain of Hec1 is not explicitly required for either Ska complex recruitment to kinetochores or for formation of stable kinetochore-microtubule attachments, but it likely plays a role in force generation at the attachment interface in human cells.

**Figure 5.**
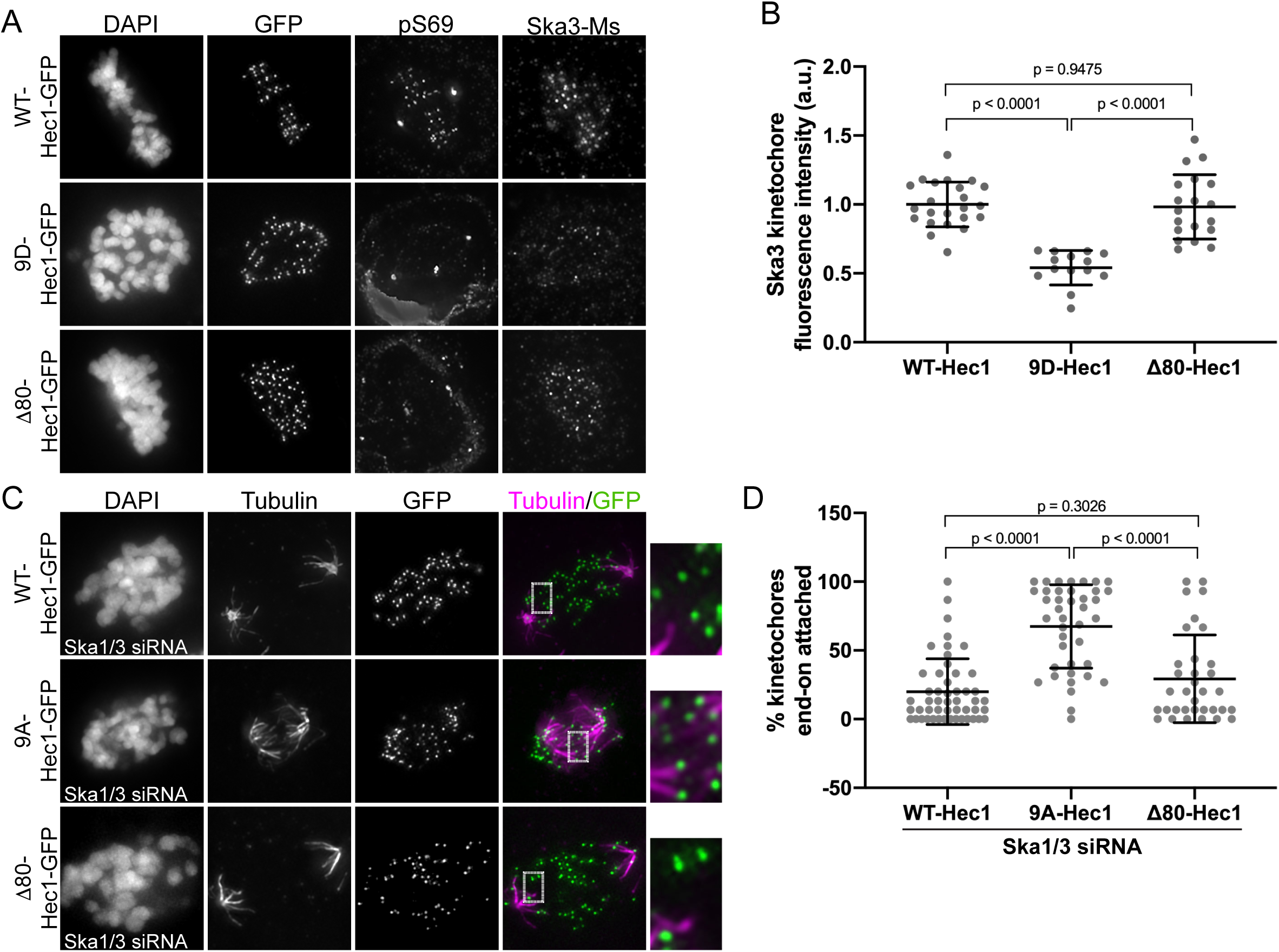
The Hec1 tail domain is dispensable for Ska complex recruitment to kinetochores, and is required for kinetochore-microtubule attachments in the absence of the Ska complex. (A) Immunofluorescence images of cells expressing WT, 9D-, and Δ80-Hec1-GFP. Cells were fixed and stained using antibodies to Hec1 pS69 and Ska3 (generated in mouse). (B) Quantification of Ska3 kinetochore fluorescence intensity from cells expressing WT, 9D-, and Δ80-Hec1-GFP. For each condition, at least 20 kinetochores per cell were measured from at least 5 cells per experiment from 3 separate experiments. Statistical significance was determined by a one-way Anova analysis. (C) Immunofluorescence images of cold-treated cells expressing WT-, 9A-, and Δ80-Hec1-GFP and treated with Ska1 and Ska3 siRNA. Cells were incubated in cold DMEM on ice for 12 minutes prior to fixation, permeabilized, fixed, and stained with antibodies to tubulin. Insets are enlargements of the region indicated by the dashed box. (D) Quantification of end-on attachment in cells expressing WT-, 9A-, and Δ80-Hec1-GFP and treated with Ska1 and Ska3 siRNA. The WT- and 9A-Hec1 data shown are from the experiment presented in Figure 2. For each condition, at least 15 kinetochores per cell were measured from at least 10 cells per experiment from 3 separate experiments. Statistical significance was determined by a one-way Anova analysis.

### The Ska complex is recruited to the internal coiled-coil domain region of the NDC80 complex to enhance NDC80-MT binding

To identify the Ska complex recruitment domain within the NDC80 complex, we carried out two color fluorescence localization mapping of Ska complex components at metaphase kinetochores (Wan et al., 2009). Since the C-terminal half of Ska3 contains the putative NDC80 binding site, we first mapped the distance between a Ska3 antibody that recognizes amino acids 226-253 (Fig. 6 A; “Ska3-M,” for “middle”) and both CENP-C (inner kinetochore), and the N-terminus of Hec1 (outer kinetochore). These measurements revealed that amino acids 226-253 of Ska3 reside ∼47 nm outside of CENP-C and ∼ 29 nm inside of the N-terminus of Hec1 (Fig 6 A-D), suggesting that a region encompassed by the NDC80 complex-binding domain is localized near the internal, coiled-coil region of the NDC80 complex. Reconstituted, purified Ska complexes have been shown to exist as either monomers or dimers of the Ska1, Ska2, and Ska3 trimer, which are formed through oligomerization of the N-termini of each protein to form a three-helix bundle (Jeyaprakash et al., 2012; Helgeson et al., 2018). Ska1’s C-terminus contains a winged-helix domain that has microtubule binding activity, and Ska3 contains a predominantly unstructured C-terminal region that is responsible for interaction with the NDC80 complex (Jeyaprakash et al., 2012; Abad et al., 2014; Abad et al., 2016). To better understand how the Ska complex components are organized at the kinetochore-microtubule interface, we carried out further paired fluorescence localization mapping using N- and C-terminal GFP tags on the Ska complex components. The N-terminal GFP tags on Ska1, Ska2, and Ska3 all mapped to a similar domain within the kinetochore which was between 52 and 65 nm outside of CENP-C and between 12 and 24 nm inside the CH domain of Hec1 (Fig. 6 C and D). This is not surprising, since the N-termini of Ska1, 2, and 3 form a well-folded, relatively compact oligomerization domain (Jeyaprakash et al., 2012). Furthermore, we found that all C-terminal domains of Ska1, Ska2, and Ska3 also mapped to a region inside of the Hec1 CH domain. However, we note the C-terminal GFP tag on Ska3 was localized very close to this region, with a mapped distance of ∼73 nm outside of CENP-C and ∼6 nm inside of the Hec1 CH domain (Fig. 6 B-D). This suggests that the unstructured domain of Ska3 may extend substantially along the length of the coiled-coil domain of the NDC80 complex. These experiments were carried out using a 2D analysis of kinetochore domain localization (Suzuki et al., 2018), and we note our reported average distance between CENP-C and the CH domain of Hec1 is consistent with previously reported 2D and 3D measurements (Fig. 6 C and D; Wan et al., 2009; Suzuki et al., 2018).

**Figure 6.**
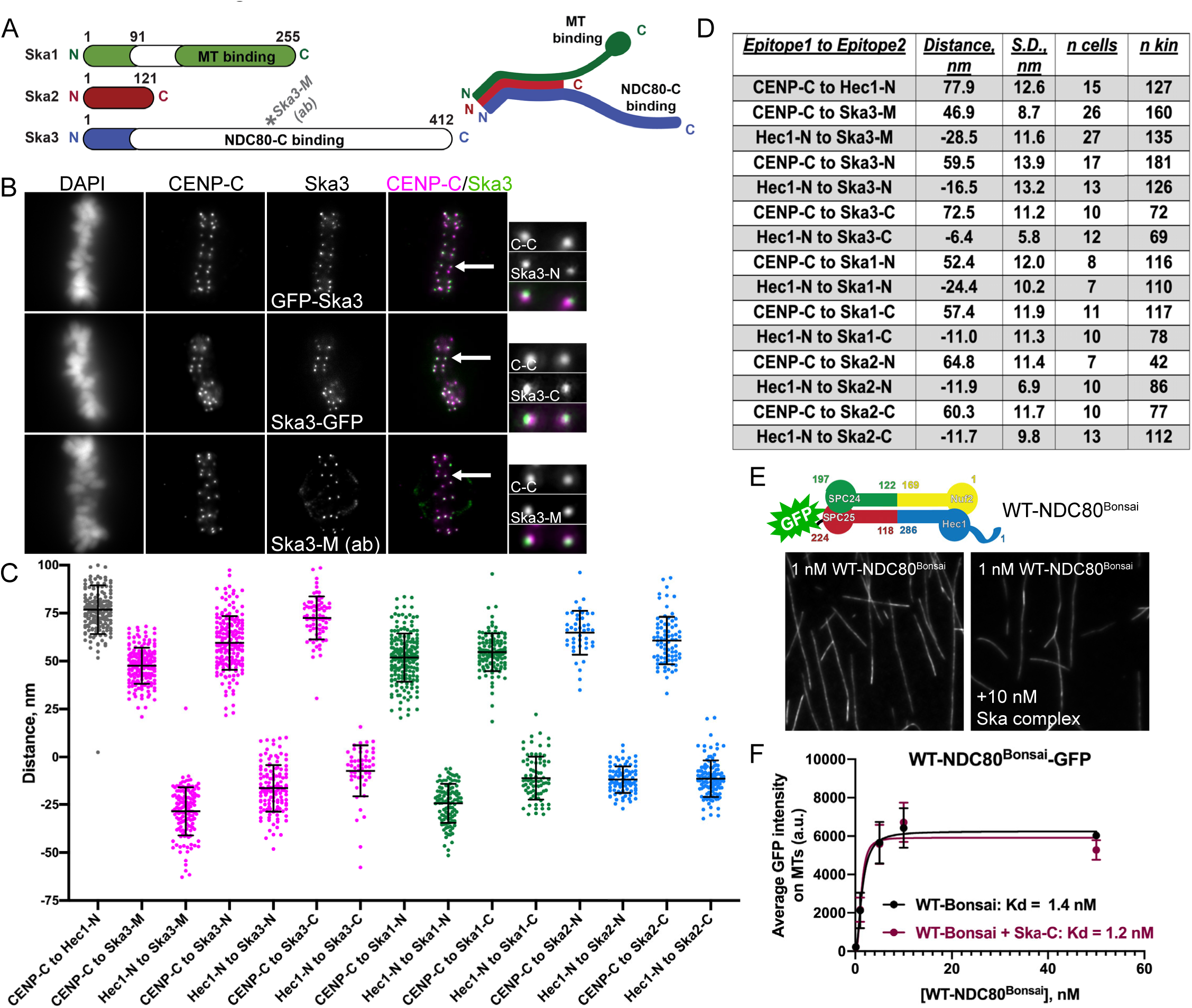
The Ska complex localizes to kinetochores at the central coiled-coil domain of the NDC80 complex. (A) Left: Schematic showing the domain architecture of the Ska complex components. White regions indicate predicted disordered domains (Jeyaprakash et al., 2012). The Ska3 antibody directed to amino acids 226-253 is indicated on the schematic and represents “Ska3-Middle Domain”, or Ska3-M. Right: Cartoon of a single Ska complex (one copy of each subunit), showing the trimerization domains located in the N-termini of Ska1, Ska2, and Ska3, the microtubule binding domain of Ska1 (green), and the proposed NDC80 complex-binding region in Ska3 (blue). (B) Immunofluorescence images of metaphase cells expressing N- and C-terminally GFP-tagged Ska3 and stained with antibodies to inner kinetochore protein CENP-C (top two rows) and immunofluorescence images of a metaphase cell stained with antibodies to Ska3-M (rabbit) and CENP-C (bottom row). Arrows point to the kinetochore pairs shown in the insets. (C) Plots of the mean distance between the indicated kinetochore proteins/protein domains. Measurements with “Hec1-N” were carried out with an antibody to the CH domain in the N-terminus of Hec1 (9G3). “N” and “C” epitopes for each of the Ska complex components are N- and C-terminal GFP moieties, respectively. Each point on the graph represents a distance measurement for a pair of sister kinetochores. (D) Summary of data presented in (C). Positive values indicate that epitope 1 was mapped inside of epitope 2. Negative values indicate that epitope 1 was mapped outside of epitope 2. S.D. indicates standard deviation. The number of cells (n cells) and kinetochore pairs (n kin) are indicated. (E) Top: Schematic of the NDC80^Bonsai^ complex. Bottom: GFP fluorescence images of the NDC80^Bonsai^ complex decorating microtubules in the presence and absence of Ska complex. Images show a single concentration of the NDC80^Bonsai^ complex from the experiment (1 nM) with and without added Ska complex (10 nM). (F) Binding curves from the microtubule binding assays. Datapoints and curve fits shown in black are from experiments without added Ska complex. Those shown in burgundy are from experiments with added Ska complex. Each point on the curve represents the average fluorescence intensity at that concentration from three separate experiments. For each concentration, fluorescence intensities of GFP-NDC80 complexes were measured on at least 40 individual microtubules from at least 10 different TIRF fields per experiment.

The mapping experiments suggested that the Ska complex is recruited to the central coiled-coil region of the NDC80 complex. To further investigate a role for this region for Ska complex binding, we carried out microtubule binding experiments in the presence and absence of purified Ska complexes using NDC80^Bonsai^ constructs, which encode for truncated NDC80 complexes missing most of the central coiled-coil region and the loop domain (Ciferri et al., 2008). Indeed, we found that while the NDC80^Bonsai^ complexes bound robustly to microtubules, the affinity of NDC80^Bonsai^ complexes for microtubules was not increased with the addition of Ska complexes (Fig. 6 E and F). These findings are consistent with recent results from the Liu and Musacchio labs, which demonstrate that NDC80^Bonsai^ complexes are unable to bind to purified Ska complexes (Zhang et al., 2017; Huis in’t Veld et al., 2019). Additionally, we found that in BRB80 buffer, addition of the Ska complex did not induce clustering of NDC80^Bonsai^ complexes on microtubules, in contrast to NDC80^Bronsai^ complexes (Fig S3 B), further supporting the idea that the central coiled-coil region of the NDC80 complex is required for Ska complex association.

We next generated a version of NDC80^Bronsai^ in which the amino acids that make up the “loop” region of Hec1 (amino acids 420-460; Maiolica et al., 2007) were substituted with alternative amino acids predicted to form a flexible motif (Varma et al., 2012) (ML-NDC80^Bronsai^; Fig. 7 A). We then tested if the microtubule binding affinity of this mutant version of the complex was increased by addition of purified Ska complexes. We found that ML-NDC80^Bronsai^ bound to microtubules with similar affinity to WT-NDC80^Bronsai^, but the addition of purified Ska complex had no significant effect on its microtubule binding affinity (Fig. 7 B and C). These results suggest that mutation of the loop domain either prevents the Ska complex from directly interacting with the NDC80 complex or precludes a conformation that promotes formation of a NDC80/Ska/microtubule complex.

**Figure 7.**
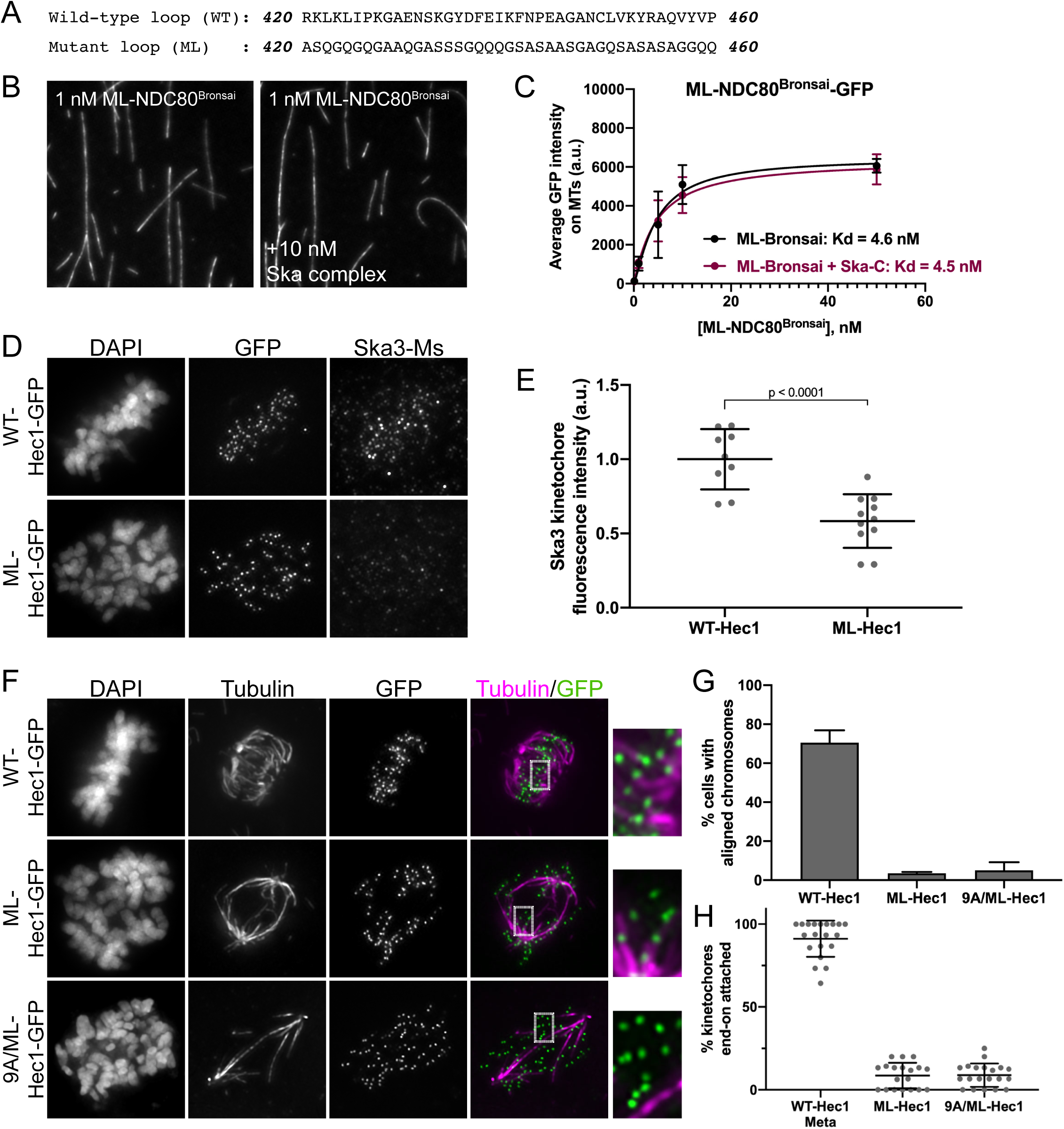
The Hec1 loop domain contributes to Ska complex recruitment to kinetochores and generation of kinetochore-microtubule attachments. (A) Sequence of the wild-type (WT) and mutated (ML) loop region in Hec1. (B) GFP fluorescence images of the ML-NDC80^Bronsai^ complex decorating microtubules in the presence and absence of the Ska complex. Images show a single concentration of the ML-NDC80^Bronsai^ complex from the experiment (1 nM) with and without added Ska complex (10 nM). (C) Binding curves from the microtubule binding assays. Datapoints and curve fits shown in black are from experiments without added Ska complex. Those shown in burgundy are from experiments with added Ska complex. Each point on the curve represents the average fluorescence intensity at that concentration from three separate experiments. For each concentration, fluorescence intensities of GFP-NDC80 complexes were measured on at least 40 individual microtubules from at least 10 different TIRF fields per experiment. (D) Immunofluorescence images of cells expressing WT- and ML-Hec1-GFP. Cells were fixed and stained using antibodies to Ska3 (mouse). (E) Quantification of Ska3 kinetochore fluorescence intensity from cells expressing WT- and ML-Hec1-GFP. For each condition, at least 20 kinetochores per cell were measured from at least 4 cells per experiment from 2 separate experiments. (F) Immunofluorescence images of cold-treated cells expressing WT-, ML-, and 9A/ML-Hec1-GFP. Cells were incubated in cold DMEM on ice for 12 minutes prior to fixation, permeabilized, fixed, and stained with antibodies to tubulin. Insets are enlargements of the region indicated by the dashed box. (G) Quantification of chromosome alignment in cells expressing WT-, ML-, and 9A/ML-Hec1-GFP. For each condition, chromosome alignment was assessed in at least 100 cells per experiments from 2 separate experiments. Cells were scored as “aligned” if they contained a metaphase plate with <5 chromosomes off the plate. (H) Quantification of end-on attachment in cells expressing WT-, ML-, and 9A/ML-Hec1-GFP. For each condition, at least 15 kinetochores per cell were measured from at least 9 cells per experiment from 2 separate experiments.

We then asked if the loop domain was required for Ska complex recruitment to kinetochores in cells. For this purpose, we expressed a version of the ML-Hec1-GFP mutant in HeLa cells, and found that Ska3 levels were significantly reduced at kinetochores compared to kinetochores from cells expressing WT-Hec1-GFP (Fig. 7 D and E). We also found, consistent with previously published results, that end-on kinetochore-microtubule attachments failed to form and chromosome alignment was abolished in cells expressing ML-Hec1-GFP (Fig. 7 F-H) (Varma et al., 2012; Zhang et al., 2012). Given that Ska complexes preferentially load to kinetochores with end-on attachments, again we could not distinguish between two possibilities: (1) the Hec1 loop domain promotes Ska complex recruitment, and in turn, the Ska complex is required for end-on attachment formation; or (2) the Hec1 loop domain is required for generation of stable kinetochore-microtubule attachments, and in turn, stable attachments promote Ska complex loading. We therefore measured Ska complex loading to kinetochores in the absence of microtubules, and found that cells expressing ML-Hec1-GFP exhibited reduced levels of Ska3 at kinetochores (Fig. 2 C and D), suggesting that an in-tact loop domain is required for efficient Ska complex recruitment to kinetochores.

Both chromosome alignment and kinetochore-microtubule attachment formation were severely impaired in cells expressing ML-Hec1-GFP. We therefore tested if these defects were exclusively due to loss of Ska complex recruitment to kinetochores. Experiments in Figure 2 demonstrated that cells expressing 9A-Hec1-GFP formed hyper-stable kinetochore-microtubule attachments, and this phenotype was independent of the Ska complex (Fig. 2 E and F). These results indicate that attachment defects arising from Ska1/Ska3 depletion can be compensated for by the strong attachments generated in cells expressing 9A-Hec1-GFP. We therefore reasoned that if a mutated loop domain results in attachment defects solely due to loss of Ska complex recruitment, then preventing phosphorylation of the tail domain should rescue this defect. Thus, we generated a hybrid mutant containing a 9A tail domain and the mutant loop sequence (9A/ML-Hec1-GFP). We found that in cells expressing 9A/ML-Hec1-GFP, stable kinetochore-microtubule attachments failed to form and chromosome alignment was severely defective, similar to what we observed in cells expressing ML-Hec1-GFP (Fig. 7 F-H). These results suggest that although the loop domain may participate in recruiting the Ska complex to kinetochores, it likely plays an additional, non-Ska complex-dependent role in generating kinetochore-microtubule attachments, perhaps through recruitment of other kinetochore proteins. It is also possible that mutation of the loop domain results in changes in NDC80 complex architecture at kinetochores that preclude formation of end-on, stable kinetochore-microtubule attachments.

To further investigate loop domain function, we generated systematic alanine substitutions of short stretches of 5-6 amino acids within the Hec1 loop (Fig. S6 A) and expressed these mutants in cells. Several, but not all, of these mutants mimicked the ML-Hec1-GFP mutant and led to severe chromosome misalignment (Fig. S6 A and B). We noted that mutating regions within the loop that resulted in substantial changes in local net charge produced the strongest chromosome misalignment phenotypes, while mutating regions with low net charge density resulted in no observable defects (Fig. S6 A-D). This suggests that the distributed charge of the loop region is likely critical for formation of kinetochore-microtubule attachments, potentially through forming interactions with the Ska complex and/or other kinetochore associated proteins such as Cdt1 (Varma et al., 2012).

## Discussion

### Hec1 tail phosphorylation affects kinetochore-microtubule attachments independently of the Ska complex

The positively-charged, N-terminal tail domain of Ndc80/Hec1 is a target of Aurora kinases, and it has been suggested that phosphorylation of the tail directly reduces affinity of NDC80 complexes for the negatively-charged microtubule lattice, which in turn reduces kinetochore-microtubule attachment strength. It is also possible that phosphorylation of the Hec1 tail domain indirectly affects kinetochore-microtubule attachment strength by regulating the recruitment of additional kinetochore-associated microtubule-binding proteins. One possible candidate for imparting this regulation is the Ska complex, which loads to kinetochores progressively during mitosis and contributes to the stabilization of kinetochore-microtubule attachments. We found here that cells expressing a non-phosphorylatable Hec1 mutant (9A-Hec1) recruit increased levels of Ska complex components to kinetochores in human cells, similar to what has been reported in *C. elegans* (Cheerambathur et al., 2017). However, we found that stable kinetochore-microtubule attachments formed in cells expressing 9A-Hec1 constructs were not dependent on the presence of a functional Ska complex. Furthermore, we demonstrated that the phosphorylation state of the tail domain did not affect the levels of Ska complexes recruited to kinetochores in the absence of microtubules. Collectively, these results suggest that while Ska complex enrichment at kinetochores coincides with Hec1 tail domain dephosphorylation, they each likely contribute to kinetochore-microtubule attachment stabilization independently.

### The Hec1 tail is dispensable for kinetochore-microtubule attachments in cells

A surprising finding from our study is that the N-terminal tail domain of Hec1 is not explicitly required for the formation of kinetochore-microtubule attachments in human cells. Consistent with our findings here, chromosome alignment errors and decreased inter-kinetochore distances have been previously observed in mammalian cells expressing tail-less Hec1 (Guimaraes et al., 2008; Miller et al., 2008; Etemad et al., 2015; Janczyk et al., 2017). These phenotypes have been widely attributed to lack of stable kinetochore-microtubule attachments in Δ80-Hec1 expressing cells, a result which has also been previously observed (Miller et al., 2008; Guimaraes et al., 2008). Contrary to this, we find that Δ80-Hec1 expression does not prevent formation of cold-resistant, stable kinetochore-microtubule attachments. Instead, we report that cells lacking the tail domain form kinetochore-microtubule attachments prematurely, which is presumably due to a lack of Aurora kinase-mediated regulation. In such a scenario, cells expressing tail-less Hec1 are unable to negatively regulate the initial formation of end-on kinetochore-microtubule attachments, and the NDC80 complex is able to bind spindle microtubule plus-ends through strong Hec1 CH domain-mediated interactions, which would otherwise be kept labile by a highly phosphorylated Hec1 tail domain.

Despite this early accumulation of attachments, however, we found that cells expressing Δ80-Hec1-GFP exhibited low inter-kinetochore distances, suggesting attachments are unable to produce sufficient forces to generate wild-type tension across sister kinetochore pairs. Alterations in force generation at the kinetochore-microtubule interface have been reported to change the balance of spindle forces in a manner that disrupts motor-mediated spindle pole focusing, which can lead to multipolarity (Manning and Compton, 2007; Maiato and Logarinho, 2014). Consistent with this idea, we found that cells expressing Δ80-Hec1-GFP undergo spindle pole fragmentation during mitotic progression, whereby the mitotic spindle becomes multi-polar and chromosomes lose alignment from the metaphase plate. Given the decreased inter-kinetochore distances observed in cells expressing Δ80-Hec1-GFP, we speculate that the tail is dispensable for forming attachments, but is required for sustained force generation during mitotic progression. Such a scenario would result in kinetochores that accumulate end-on microtubule attachments early in mitosis due to the absence of Aurora kinase-mediated regulation, but exhibit defects in transducing forces from the dynamics of attached microtubules in late mitosis, leading to loss of spindle bipolarity and chromosome alignment. It is noteworthy to mention that this hypothesis is supported by recent biochemical experiments reported in Huis in’t Veld et al. (2019), where the authors artificially trimerized NDC80 complexes on the surface of beads and measured the ability of NDC80 complex trimers to resist force from an optical trap. Complexes lacking the Hec1 tail – despite binding to microtubules with high affinity – detached from depolymerizing microtubules in both the presence and absence of applied force, whereas wild-type NDC80 trimers remained bound under these conditions (Huis in’t Veld et al., 2019). These results led the authors to conclude that the Hec1 tail is critical for force-coupling at the kinetochore-microtubule interface.

Our results describing the formation of end-on attachments in human cells expressing tail-less Hec1 are consistent with results from both budding yeast and *C. elegans,* where the Ndc80 tail domain is not strictly required for kinetochore-microtubule attachment (Kemmler et al., 2009; Demirel et al., 2012; Lampert et al., 2013; Cheerambathur et al., 2013). However, similar to the scenario in human cells, the tail domain does play some role at the kinetochore-microtubule interface in these organisms. For example, in budding yeast the tail domain becomes required for cell survival upon perturbation of the Dam1 complex, a kinetochore-associated complex found in yeasts but not higher eukaryotes, which contributes to generating stable kinetochore-microtubule attachments (Demirel et al., 2012; Lampert et al., 2013). In addition, a study using a tension sensor near the N-terminus of budding yeast Hec1/Ndc80 demonstrated that while the tail domain is not required for kinetochore-microtubule attachments *per se* in cells, its deletion results in loss of tension at the NDC80 complex-microtubule interface, which can be compensated for with an in-tact Dam1 complex (Suzuki et al., 2016). Taken with the result that the Hec1 tail is required for load-bearing attachments of NDC80 complexes to microtubules (Huis in’t Veld et al., 2019), the available data suggest that this function of the Hec1/Ndc80 tail domain is generally conserved from yeast to humans.

### The Ska complex compensates for Hec1 tail domain function

Studies using NDC80 complexes purified from various organisms have demonstrated that the Hec1 tail domain is required for high affinity NDC80 complex-microtubule binding *in vitro* (Cheeseman et al., 2006; Wei et al., 2007; Lampert et al., 2013; Alushin et al., 2012; Umbreit et al., 2012; Cheerambathur et al., 2013; Zaytsev et al., 2015). In the case of human NDC80 complexes, addition of the Ska complex compensates for deletion of the Hec1 tail in a number of *in vitro* NDC80 complex-microtubule interaction assays. The Davis lab carried out optical trapping experiments using NDC80 complex-coated beads to demonstrate that while NDC80 complexes lacking the Hec1 tail generated weak attachments to microtubules that could be disrupted under low rupture forces, addition of soluble Ska complexes significantly strengthened these attachments (Helgeson et al., 2018). Consistent with these findings, Huis in’t Veld et al. found that addition of the Ska complex to trimerized NDC80 complexes lacking the Hec1 tail enabled these complexes to track depolymerizing microtubules, a property not observed in the absence of the Ska complex (Huis in’t Veld et al., 2019). The notion that the Ska complex can functionally compensate for Hec1 tail deletion is reminiscent of studies carried out in budding yeast with the Dam1 complex, which has been suggested to be a functional ortholog of the Ska complex (Welburn et al., 2009). Analogous to the experiments described above for human Ska and NDC80 complexes, Dam1 is able to enhance the affinity of budding yeast NDC80 complexes for microtubules *in vitro* (Tien et al., 2010; Lampert et al., 2010; Lampert et al., 2013). Consistently, as mentioned above, deletion of the Ndc80/Hec1 tail is not lethal in budding yeast (Kemmler et al., 2009; Demirel et al., 2012; Lampert et al., 2013), but deletion or mutation of Dam1 sensitizes cells to Ndc80/Hec1 tail deletion, resulting in cell death due to cell division defects (Demirel et al., 2012; Lampert et al., 2013; Suzuki et al., 2016). In line with these results we found that cells expressing Δ80-Hec1-GFP were able to form cold-stable kinetochore-microtubule attachments, but this required the presence of the Ska complex. Collectively, our *in vitro* and cell-based results suggest that the Ska complex can compensate for the Hec1 tail’s role in forming stable kinetochore-microtubule attachments in human cells.

### The Ska complex is recruited to the internal coiled-coil domain of the NDC80 complex rather than the Hec1 tail domain

Ska complex loading to kinetochores requires the NDC80 complex (Gaitanos et al., 2009; Welburn et al., 2009; Chan et al., 2012; Zhang et al., 2012), and the two complexes directly interact (Zhang et al., 2017; Helgeson et al., 2018; Huis in’t Veld et al., 2019). Although Ska3 is known to mediate the interaction, its binding site on the NDC80 complex remains unresolved (Zhang et al., 2018). Our results demonstrate that the Hec1 tail is not required for Ska complex-mediated enhancement of NDC80-microtubule interactions or for Ska complex localization to kinetochores in cells. We note that these results are inconsistent with a previous report from the Stukenberg lab, where it was shown that mutations in the Hec1 tail abolish Ska recruitment to the NDC80 complex-microtubule interface *in vitro* and to kinetochores in cells (Janczyk et al., 2017). It is unclear why our results differ from theirs, although a potential explanation is that the tail mutations made in the Janczyk et al. study impacted overall kinetochore architecture in a manner that precluded Ska recruitment independently of the Hec1 tail. To map the Ska complex kinetochore recruitment domain, we used two-color co-localization imaging and found that the Ska complex co-localizes with the coiled-coil region of the NDC80 complex, inside of the Hec1 CH domain. Interestingly, most of the N- and C-termini of all Ska complex components also map near this region, suggesting that the bulk of the complex is not significantly extended along the NDC80 complex axis. The one exception is the C-terminus of Ska3, which maps closely to, but still inside of, the CH domain of Hec1, suggesting that the unstructured region of Ska3 may be somewhat elongated along the length of the NDC80 complex. These results are consistent with results from the Davis lab, which found a large number of contact points between Ska3 and the coiled-coil region of the NDC80 complex using a cross-linking mass spectrometry technique (Helgeson et al., 2018).

### The Hec1 loop domain has Ska complex-dependent and -independent functions in chromosome alignment

We report that mutation of the Hec1 loop domain prevents enhancement of NDC80 complex-microtubule binding by the Ska complex. This is possibly at odds with a recent study from the Musacchio lab, which reports that removal of the loop domain from Hec1 does not affect the interaction between soluble NDC80 and Ska complexes (Huis in’t Veld et al., 2019). This difference could possibly reflect a requirement for the loop domain in the interaction of NDC80 and Ska complexes specifically on microtubules. Alternatively, since Ska3 phosphorylation by CDK1 increases the affinity of soluble Ska and NDC80 complexes for each other (Zhang et al., 2017; Huis in’t Veld et al., 2019), the phosphorylation state of Ska3 might impact the requirement of the Hec1 loop domain for the two complexes to associate. In such a scenario, dephosphorylated Ska complexes would also bind more weakly to NDC80 complexes in the presence of microtubules, and the loop domain would be required for high affinity interactions specifically under these sub-optimal binding conditions. Given our result that the loop domain is required in cells for microtubule-independent Ska complex loading to kinetochores, we do not favor the latter hypothesis.

Both chromosome alignment and kinetochore-microtubule attachments were severely perturbed in cells expressing Hec1 constructs containing a mutated loop domain -- more so than in cells expressing 9D-Hec1 mutants or in cells depleted of Ska1 and Ska3. This was observed in previous studies (Zhang et al., 2012; Varma et al., 2012), and led us to ask if the defects observed were entirely due to loss of Ska complex recruitment. To test this, we modified the loop mutant to include a non-phosphorylatable N-terminal tail domain, since we demonstrated that expression of the 9A-Hec1 mutant abrogated the need for Ska1 and Ska3 to form stable kinetochore-microtubule attachments. Cells expressing the 9A/ML-Hec1 mutant showed no improvement in either chromosome alignment or formation of stable kinetochore-microtubule attachments compared to those expressing ML-Hec1. Thus, we conclude that in addition to contributing to efficient Ska complex recruitment to kinetochores, the Hec1 loop has an additional, non-Ska complex-dependent role in forming stable attachments. We found that mutating short stretches of the loop sequence that contain at least two charged residues phenocopied expression of the full ML-Hec1 construct. Thus it is possible that the loop domain recruits additional factors, such as Cdt1, that are required for generation of stable, end-on kinetochore-microtubule attachments (Varma et al., 2012). Alternatively, the loop region could be critical for adoption of a conformation of NDC80 that is required for maintaining proper, end-on attachments to microtubules.

Overall, our results support a model in which the Hec1 tail domain, while not explicitly required for forming end-on kinetochore-microtubule attachments, is important in regulating the force-generating attachments between kinetochores and microtubule plus ends. They also suggest that the Ska complex is loaded to the central coiled-coil region of the NDC80 complex during mitosis to ensure proper force-coupling at the kinetochore-microtubule interface, which may be particularly important at kinetochores containing NDC80 complexes with lower microtubule binding capacity (e.g. with highly phosphorylated tail domains). How tail domain phosphorylation and dephosphorylation are coordinated with Ska complex loading to kinetochores to ensure proper regulation of kinetochore-microtubule attachments is an important issue that begs future investigation.

## Materials and Methods

### Cell culture

HeLa Kyoto cells were cultured in Dulbecco’s Modified Eagle Medium (DMEM) supplemented with 10% FBS and 1% antibiotic/antimycotic solution. RPE-1 (ATCC) cells were cultured in 1:1 Ham’s F12:DMEM supplemented with 10% FBS and 1% antibiotic/antimycotic solution. All cell lines were maintained at 37°C in 5% CO_2_.

### Cell treatments and transfections

For all fixed-cell experiments, cells were grown on sterile, acid-washed coverslips in six well plates. All nucleic acid transfections were done in Optimem (Gibco). siRNA duplexes were transfected as follows: Ska3 (5′-AGACAAACAUGAACAUUAA-3′; Gaiatanos et al., 2009) at 80 nM, Ska1 (5′-CCCGCTTAACCTATAATCAAA-3′; Hanisch et al., 2006) at 80 nM, and Hec1 (5′-CCCUGGGUCGUGUCAGGAA-3′; DeLuca et al., 2011) at 160 nM. All siRNA duplexes were transfected into HeLa cells using Oligofectamine (Thermo Fisher Scientific) according to the manufacturer’s protocol. For all siRNA transfections, cells were processed for immunofluorescence 48 hours after addition of siRNA. Plasmids encoding Hec1-GFP (2 µg) were transfected into HeLa Kyoto cells with Lipofectamine 2000 (Thermo Fisher Scientific) according to the manufacturer’s protocol. RPE1 cells were transfected using a nucleofector (Lonza) and using Lonza Kit L according to the manufacturer’s protocol, with program X-001. 24 hours post-DNA transfection, cells were processed for immunofluorescence. For silence-rescue experiments of Hec1 and expression of Hec1 mutants in Ska-depleted cells, cells were transfected with siRNA using Oligofectamine, and 24 hours later were transfected with DNA using Lipofectamine 2000. Twenty-four hours after DNA transfection (48 hours post-siRNA addition), cells were processed for immunofluorescence.

For end-on attachment experiments, transfection media was replaced with ice cold DMEM 24 hours post-DNA transfection and cells were incubated on ice for 12 minutes prior to fixation. For attachment-independent analysis of Ska3 localization, Hec1-GFP-transfected cells were arrested at the G2/M transition with 9 µM RO3306 (Sigma-Aldrich) for 16 hours, then extensively washed out into 10 µM nocodazole (Tocris Bioscience) for one hour before fixation and subsequent processing for immunofluorescence. For analysis of asynchronous attachment-independent Ska3 levels (Figure S2), cycling HeLa cell populations were treated with 10 µM nocodazole for one hour, then fixed.

### Immunofluorescence Microscopy

Cells were rinsed in PHEM buffer (60 mM PIPES, 25 mM HEPES, 10 mM EGTA, 4 mM MgCl_2_, pH 7.0) and permeabilized in lysis buffer (PHEM + 1.0% Triton-X 100) for 5 minutes at 37°C. Post-lysis, cells were quickly washed in PHEM and subsequently fixed in freshly made fixative solution (4% paraformaldehyde in PHEM) for 20 minutes at 37°C. After fixation, cells were subjected to three five-minute washes in PHEM-T (PHEM + 0.1% Triton-X 100), quickly rinsed in PHEM, and blocked in 10% boiled donkey serum (BDS) in PHEM for 1 hour at room temperature. Following blocking, primary antibodies diluted in 5% BDS in PHEM were added to cells and incubated for 1 hour at room temperature followed by 16 hours at 4°C. Primary antibodies were used as follows: human anti-centromere antibody (ACA) at 1:300 (Antibodies, Inc.), mouse anti-tubulin (DM1ɑ) at 1:600 (Sigma-Aldrich), mouse anti-Hec1 (9G3) at 1:3000 (Novus Biologicals), rabbit anti-phosphorylated Hec1-pSer69 (pS69) at 1:3000 (DeLuca et al., 2018), mouse anti-Ska3 at 1:500 (Santa-Cruz Biotechnology), rabbit anti-Ska3 at 1:300 (GeneTex), and rabbit anti-Astrin at 1:1000 (Sigma-Aldrich). After primary antibody incubation, unbound antibody was washed off using three five-minute PHEM-T rinses, followed by a quick wash in PHEM. Secondary antibodies (conjugated to Alexa 488, Cy3 dye, or Alexa 647, Jackson ImmunoResearch) were diluted 1:1000 in 5% BDS except mouse Ska3 and rabbit Ska3, for which secondary antibodies were diluted 1:500 and 1:300, respectively. Cells were incubated in secondary antibody for 45 minutes at room temperature, and unbound antibody was washed off with 3 x 5 min PHEM-T washes followed by a quick rinse in PHEM. Cells were then incubated in a 2 ng/ml DAPI solution (diluted in PHEM) for 30 seconds, subjected to two five-minute PHEM-T washes, quickly rinsed in PHEM, and mounted onto glass slides using an antifade solution (90% glycerol + 0.5% *N*-propyl gallate). Following mounting, coverslip edges were sealed with nail polish and slides were stored at 4°C.

### Fixed cell imaging

All fixed cell images were acquired using a DeltaVision Personal DV Imaging system (GE Healthcare) on an IX71 inverted microscope (Olympus) using SoftWoRx software (GE Healthcare). All fixed cell experiments were imaged using a 60X 1.42 NA differential interference contrast Plan Achromat oil immersion lens (Olympus). Images were acquired using a CoolSNAP HQ2 camera (Photometrics/Roper Technologies) for a final magnification of 107 nm/pixel. For two-color distance measurements, a 1.6 magnification lens was inserted in the light path, providing a final magnification of 67 nm/pixel at the camera sensor.

### Live cell imaging

For live cell imaging experiments, cells were seeded into custom built glass-bottom 35 mm dishes and imaged in Leibovitz’s L-15 medium (Invitrogen) supplemented with 10% FBS, 7 mM Hepes, and 4.5 g/liter glucose, pH 7.0. For live cell experiments, HeLa Kyoto cells were transfected with plasmids encoding Hec1-GFP (2 µg) and mCherry-tubulin (300 ng) as described above. 24 hours post-transfection, cells were imaged on a Nikon Ti-E microscope equipped with a Piezo Z-control (Physik Instrumente), stage top incubation system (Okolab), spinning disc confocal scanner unit (CSUX1; Yokogawa) using a 1.49 NA oil immersion 100X objective and an iXon DU888 EM-CCD camera (Andor) for a final magnification of 130nm/pixel. Z-stacks were acquired taking 7 planes at 1 µm steps using 488 nm and 594 nm lasers to excite GFP and mCherry, respectively. Transfection-positive bipolar cells were identified and imaged for 3 hours using 3 and 5 minute intervals.

### Protein expression and purification

Glutathione-S-transferase (GST)-NDC80^Bonsai^ (Ciferri et al., 2008) was a generous gift from Andrea Musacchio (Max Planck Institute of Molecular Physiology, Dortmund, Germany). GST-NDC80^Bronsai^ constructs were generated from GST-NDC80^Bonsai^ (parent vector backbone = pGEX6P1-2RBS). Specifically, Nuf2^1-348^/Spc24^122-197^, Hec1^1-506^/Spc25^118-224^ fragments were obtained by PCR from parent vectors of each protein while creating 20bp overhangs for Gibson reaction for cloning back into BamH1/Age1 digested GST-Bonsai plasmid. Cloning of Δ80-NDC80^Bronsai^ was carried out using the same fragments, except the Hec1 PCR used a forward primer with amino acid 81 immediately following the start codon. ML-NDC80^Bronsai^ was generated by producing a PCR fragment of Hec1^1-461^ with the mutant loop sequence from the cell expression vector used in this study, and annealing it into the NDC80^Bronsai^ vector digested with Sac1/Afl2. NDC80^Bonsai^ and NDC80^Bronsai^ constructs were expressed and purified using the following scheme: BL21-DE3 cells were transformed with NDC80 constructs, and cultures were grown to the appropriate OD_600_ before induction overnight (16 hours) at 18°C with 400 µM isopropyl β-D-1-thiogalactopyranoside. All steps after induction were carried out at 4°C. The next morning, cells were pelleted by centrifugation and resuspended in lysis buffer (25 mM Tris pH 7.6, 300 mM NaCl, 1 mM EDTA) supplemented with protease inhibitors (Pierce Protease Inhibitor tablets, Thermo Scientific) and 1 mM dithiothreitol (DTT, Gold Bio). Resuspended cells were lysed using a microfluidic chamber at 80 psi. The resulting lysed mixture was cleared of cell debris by centrifugation at 40,000 rpm for 45 minutes in a Beckman L8-70M ultracentrifuge using a TY70-TI rotor. Supernatant was applied to glutathione-agarose resin (Pierce resin, Thermo Scientific) (pre-equilibrated in lysis buffer), and the mixture was rocked gently for one hour. Following binding, unbound protein was washed from the resin with lysis buffer, and resin-bound protein was eluted by GST-tag cleavage overnight with Human Rhinovirus 3C Protease (HRV3C Protease, expressed and purified in-house). Elutions were pooled, concentrated, and run on a GE Superdex 200 HiLoad 16/60 sizing column in lysis buffer supplemented with 5% glycerol and 1 mM DTT. Protein fractions were pooled and concentrated, and glycerol was added to 20% final volume before small aliquots were snap-frozen in liquid nitrogen and stored at −80°C.

Purification of recombinant human Ska complex (SkaC) was carried out as described previously (Abad et al., 2016). Briefly, BL21-Gold E. coli cells were co-transformed with equal amounts of the individual Ska1, GST-Ska2, and Ska3 plasmids. Cells were grown to the appropriate OD_600_ before induction overnight (16 hours) at 18°C with 400 µM isopropyl β-D-1-thiogalactopyranoside. All steps after induction were carried out at 4°C. The next morning, cells were pelleted by centrifugation and resuspended in SkaC lysis buffer (20 mM Tris pH 8.0, 500 mM NaCl) supplemented with protease inhibitors and 5 mM DTT. Resuspended cells were lysed by microfluidics as specified in NDC80 complex purifications. The resulting lysed mixture was cleared of cell debris by centrifugation at 40,000 rpm for 45 minutes as noted for NDC80 complex purifications. The supernatant from lysed cells was applied to glutathione-agarose resin (pre-equilibrated in SkaC lysis buffer) and rocked gently for 3 hours. Following binding, unbound protein was washed away with lysis buffer, and resin was further washed with chaperone buffer (20 mM Tris pH 8.0, 1M NaCl, 50 mM KCl, 10 mM MgCl_2_, 2 mM ATP, 5 mM DTT) to remove associated protein chaperones. Resin-bound protein was then eluted using 3 sequential elutions for one hour each in elution buffer (20 mM Tris pH 8.0, 100 mM NaCl, 50 mM Glutathione, 5 mM DTT). Elutions were pooled and dialyzed overnight into column buffer (20 mM Tris pH 8.0, 100 mM NaCl, 5 mM DTT) and tags were simultaneously cleaved with tobacco etch virus (TEV) protease overnight while rocking. Cleaved SkaC was further purified by gel filtration on a Superose 6 Increase 10/300 in column buffer. Protein-containing fractions were collected and concentrated, and protein was snap-frozen in liquid nitrogen and stored at −80°C.

### TIRF Microscopy

Immediately prior to microtubule binding assays, protein was flash-thawed and centrifuged at 90,000 × g to remove large aggregates. Supernatant was collected and concentration measured by Bradford assay. TIRF microscopy (TIRFM) binding assays were performed as described previously (Ecklund et al., 2017). Briefly, flow-chambers were constructed by adhering plasma cleaned, silanized coverslips (22×30mm) to glass slides with double-sided tape. Silanized coverslips were incubated with a rat anti-tubulin antibody (8 μg/ml, YL1/2; Accurate Chemical & Scientific Corporation) for 5 minutes, then blocked with 1% Pluronic F-127 solution (Fisher Scientific) for 5 minutes. Taxol-stabilized, Alexa647-labeled microtubules (made by mixing fluorescently labeled and unlabeled porcine tubulin at a 1:12.5 ratio) diluted in BRB80 (80 mM PIPES, 1 mM EGTA, 1 mM MgCl2) supplemented with 20 µM taxol were flowed into the chamber and incubated for 5-10 minutes, then unbound microtubules were washed out with one chamber volume of SN (“Ska-NDC80”) buffer (20 mM Tris pH 7.0, 50 mM NaCl, 6 mg/ml bovine serum albumin, 4 mM DTT, 20 µM taxol). GFP-NDC80 complex (either alone or supplemented with 10 nM unlabeled Ska complex) diluted to the appropriate concentration in SN buffer was introduced to the chamber and the binding reaction was incubated for 2 minutes. Two more additions of NDC80 complex (or NDC80 complex + Ska complex) were subsequently perfused into the chamber to allow binding reaction to reach equilibrium, and two minutes after the third addition (after binding reaction had reached equilibrium as determined by time-lapse imaging), and TIRF images were collected of 10 individual fields. For analysis in BRB80 and BRB20 (Supplemental Figure 4), all steps were performed as above, except protein (either GFP-NDC80 alone or supplemented with SkaC, or SkaC-GFP) was either diluted into BRB80 or BRB20 supplemented with 6mg/ml BSA, 4 mM DTT, and 20 µM taxol. All TIRFM images were collected at room temperature using a 1.49 NA 100 X Plan Apo TIRF oil immersion lens on a Nikon Ti-E inverted microscope equipped with an iXon3 DU897 EM-CCD camera (Andor) for a final pixel size of 160 nm/pixel.

### Data analysis

Measurement of kinetochore fluorescence intensity in fixed cells was measured from non-deconvolved, non-compressed images using a custom program in MatLab (Mathworks) courtesy of X. Wan (Wan et al., 2009). For analysis of Ska3 levels at attached kinetochores (Figure 1A-C, Figure 4G-H, Supplemental Figure 1), cells with greater than 25% Hec1-pS69 levels measured in WT-Hec1-GFP cells were discarded from analysis to reduce effects of endogenous, non-mutant kinetochore Hec1. Measurements of end-on attachment and inter-kinetochore distances in cold-treated cells were performed in SoftWoRx Explorer software. End-on attachment was analyzed by selecting random kinetochores in the kinetochore channel, then subsequently overlaying the tubulin channel and scoring whether spindle microtubules terminated at the pre-selected kinetochores (lateral attachments were not quantified). For analysis of end-on attachment in Ska-depleted cells, only kinetochores between the spindle poles were analyzed, as polar chromosomes remained unattached in all conditions (Figure 2E). Inter-kinetochore distances were analyzed by measuring the distance between Hec1-GFP signals from two kinetochores in a sister pair in the same z-plane. For chromosome alignment analysis, bipolar Hec1-GFP expressing cells post-nuclear envelope breakdown were scored as either aligned (metaphase plate with <5 chromosomes off the plate) or unaligned (no metaphase plate, or metaphase plate with 5 or more chromosomes off the plate). For analysis of multipolarity, Hec1-GFP expressing cells were stained with anti-tubulin antibodies and assessed for number of spindle poles.

Two-color super-resolution distance measurements were carried out on kinetochore pairs that resided in a single focal plane from non-deconvolved images. The centroids of each kinetochore test antibody signal were determined using SpeckleTracker (Wan et al., 2009). Distances were then calculated using the identified centroids.

For analysis of TIRFM microtubule binding assays, GFP-NDC80 complex-microtubule binding was quantitated using ImageJ software (National Institute of Health). NDC80 fluorescent signal was measured along the microtubule axis (as determined from Alexa647-tubulin signal), and “background” signal was measured using the same mask (created along the microtubule’s length) in a region immediately adjacent to the microtubule. Corrected signal intensity was measured by subtracting the background signal from the GFP signal on the microtubule. Raw GFP-NDC80 fluorescence intensity at each concentration (averaged across all 3 replicates) was plotted and curves were fitted using a Specific binding model with a Hill fit in Prism (Graphpad). All statistical analyses were carried out using Prism (Graphpad).

## Acknowledgements

We thank members of the DeLuca laboratory for helpful discussions, Drs. Ted Salmon and Xiaohu Wan for providing MatLab code to carry out kinetochore fluorescence intensity analysis and two-color fluorescence mapping experiments and Dr. Gary Gorbsky for the Ska1-GFP expression plasmid. This work was supported by National Institutes of Health grants R01GM088371 and R35GM130365 (to J.G. DeLuca).

## Author Contributions

JGD and RW conceived the project and JGD supervised the project. RW carried out the experiments and analyzed all data except for the two-color fluorescence localization experiments, which were carried out and analyzed by KFD. JEM cloned, expressed, and purified NDC80 constructs and complexes. IJS and AAJ cloned, expressed, and purified Ska complexes. JH helped with data analysis. JGD and RW wrote the paper with input from KFD and AAJ.

## References

Alushin, G. M., Musinipally, V., Matson, D., Tooley, J., Stukenberg, P. T., and Nogales, E. 2012. Multimodal microtubule binding by the Ndc80 kinetochore complex. Nature Structural & Molecular Biology, 19:1161–7.

Alushin, G. M., Ramey, V. H., Pasqualato, S., Ball, D. A., Grigorieff, N., Musacchio, A., and Nogales, E. 2010. The Ndc80 kinetochore complex forms oligomeric arrays along microtubules. Nature, 467805-10.

Auckland, P., Clarke, N. I., Royle, S. J., and McAinsh, A. D. 2017. Congressing kinetochores progressively load Ska complexes to prevent force-dependent detachment. Journal of Cell Biology, 216:1623–1639.

Bakhoum, S.F., SL. Thompson, AL. Manning, and DA Compton. 2009. Genome stability is ensured by temporal control of kinetochore-microtubule dynamics. Nat. Cell Biol. 11:27–35.

Biggins, S., FF. Severin, N. Bhalla, I. Sassoon, AA. Hyman, and AW. Murray. 1999. The conserved protein kinase Ipl1 regulates microtubule binding to kinetochores in budding yeast. Genes Dev. 13:532–44.

Carmena, M., M. Wheelock, H. Funabiki, and WC. Earnshaw. 2012. The Chromosomal Passenger Complex (CPC): from easy rider to the godfather of mitosis. Nat. Rev. Mol. Cell Biol. 13:789–803.

Chan, Y. W., Jeyaprakash, A. A., Nigg, E. A., and Santamaria, A. 2012. Aurora B controls kinetochore-microtubule attachments by inhibiting Ska complex-KMN network interaction. Journal of Cell Biology, 196:563–571.

Cheerambathur, D. K., Prevo, B., Hattersley, N., Lewellyn, L., Corbett, K. D., Oegema, K., and Desai, A. 2017. Dephosphorylation of the Ndc80 Tail Stabilizes Kinetochore-Microtubule Attachments via the Ska Complex. Developmental Cell, 41:424–437.e.4.

Cheerambathur, D. K., Gassmann, R., Cook, B., Oegema, K., and Desai, A. 2013. Crosstalk between microtubule attachment complexes ensures accurate chromosome segregation. Science, 342:1239–1242.

Cheeseman, I.M., JS. Chappie, EM. Wilson-Kubalek, and A. Desai. 2006. The conserved KMN network constitutes the core microtubule-binding site of the kinetochore. Cell. 127:983–97.

Ciferri, C., DeLuca, J. G., Monzani, S., Ferrari, K. J., Ristic, D., Wyman, C., Stark, H., Kilmartin, J., Salmon, E. D., and Musacchio, A. 2005. Architecture of the Human Ndc80-Hec1 Complex, a Critical Constituent of the Outer Kinetochore. Journal of Biological Chemistry, 280:29088–29095.

Ciferri, C., Pasqualato, S., Screpanti, E., Varetti, G., Santaguida, S., Dos Reis, G., Maiolica, A., Polka, J., DeLuca, J. G., De Wulf, P., Salek, M., Rappsilber, J., Moores, C. A., Salmon, E. D., and Musacchio, A. 2008. Implications for Kinetochore-Microtubule Attachment from the Structure of an Engineered Ndc80 Complex. Cell, 133:427–439.

Cimini, D., X. Wan, CB. Hirel, and ED. Salmon. 2006. Aurora kinase promotes turnover of kinetochore microtubules to reduce chromosome segregation errors. Curr Biol. 16:1711–1718.

Daum, J. R., Wren, J. D., Daniel, J. J., Sivakumar, S., Mcavoy, J. N., Potapova, T. A., and Gorbsky, G. J. 2009. Ska3 is Required for Spindle Checkpoint Silencing and the Maintenance of Chromosome Cohesion in Mitosis. Curr Biol, 19:1467–1472.

DeLuca, J.G., WE. Gall, C. Ciferri, D. Cimini, A. Musacchio, and ED. Salmon. 2006. Kinetochore microtubule dynamics and attachment stability are regulated by Hec1. Cell. 127:969–982.

DeLuca, J. G., and Musacchio, A. 2012. Structural organization of the kinetochore-microtubule interface. Current Opinion in Cell Biology, 24:48–56.

DeLuca, K. F., Lens, S. M., and DeLuca, J. G. 2011. Temporal changes in Hec1 phosphorylation control kinetochore-microtubule attachment stability during mitosis. J Cell Sci. 124:622–634.

DeLuca, K. F., Meppelink, A., Broad, A. J., Mick, J. E., Peersen, O. B., Petkas, S., Lens, S. M. A., and DeLuca, J. G. 2018. Aurora A kinase phosphorylates Hec1 to regulate metaphase kinetochore-microtubule dynamics. Journal of Cell Biology, 217:163–177.

Demirel, P. B., Keyes, B. E., Chatterjee, M., Remington, C. E., and Burke, D. J. 2012. A Redundant Function for the N-Terminal Tail of Ndc80 in Kinetochore-Microtubule Interaction in Saccharomyces cerevisiae. Genetics, 192:753–756.

Ecklund, K.H., Morisaki, T., Lammers, L.G., Marzo, M.G., Stasevich, T.J., and Markus, S.M. 2017. She1 affects dynein through direct interactions with the microtubule and the dynein microtubule-binding domain. Nat Commun. 8:2151.

Etemad, B., TE. Kuijt, and GJ. Kops. 2015. Kinetochore-microtubule attachment is sufficient to satisfy the human spindle assembly checkpoint. Nat. Commun. 6:8987.

Gaiatanos, T. N., Santamaria, A., Jeyaprakash, A. A., Wang, G., Conti, E., and Nigg, E. A. 2009. Stable kinetochore-microtubule interactions depend on the Ska complex and its new component Ska3/C13Orf3. The EMBO Journal, 28:1375–1377.

Guimaraes, G. J., and DeLuca, J. G. 2009. Connecting with Ska, a key complex at the kinetochore-microtubule interface. The EMBO Journal, 28:1442–1452.

Guimaraes, G. J., Dong, Y., McEwen, B. F., and DeLuca, J. G. 2008. Kinetochore-Microtubule Attachment Relies on the Disordered N-Terminal Tail Domain of Hec1. Curr Biol, 18:1778–1784.

Hanisch, A., Sillje, H. H. W., and Nigg, E. A. 2006. Timely anaphase onset requires a novel spindle and kinetochore complex comprising Ska1 and Ska2. The EMBO Journal, 25:5504–5515.

Helgeson, L. A., Zelter A., Riffle, M., Maccoss, M. J., Asbury, C. L., and Davis, T. N. 2018. Human Ska complex and Ndc80 complex interact to form a load-bearing assembly that strengthens kinetochore-microtubule attachments. PNAS, 115:2740–2745.

Huis in’t Veld, P. J., Volkov, V. A., Stender, I., Musacchio, A., and Dogterom, M. 2019. Molecular determinants of the Ska-Ndc80 interaction and their influence on microtubule tracking and force-coupling. bioRxiv preprint first posted online Jun. 20, 2019; doi: http://dx.doi.org/10.1101/675363. https://www.biorxiv.org/content/biorxiv/early/2019/06/20/675363.full.pdf

Janczyk, P., Skorupka, K. A., Tooley, J. G., Matson, D. R., Kestner, C. A., West, T., Pornillos, O., and Stukenberg, P. T. 2017. Mechanism of Ska Recruitment by Ndc80 Complexes to Kinetochores. Developmental Cell, 41:438–449.e4.

Jeyaprakash, A. A., Santamaria, A., Jayachandran, U., Chan, Y. W., Benda, C., Nigg, E. A., and Conti, E. 2012. Sructural and Functional Organization of the Ska Complex, a Key Component of the Kinetochore-Microtubule Interface. Molecular Cell, 46:274–286.

Kemmler, S., Stach, M., Knapp, M., Ortiz, J., Pfannstiel, J., Ruppert, T., and Lechner, J. 2009. Mimicking Ndc80 phosphorylation triggers spindle assembly checkpoint signaling. The EMBO Journal, 28:1099–1110.

Kettenbach, A. N., Schweppe, D. K., Faherty, B. K., Pechenick, D., Pletnev, A. A., and Gerber, S. 2011. Quantitative Phosphoproteomics Identifies Subtrates and Functional Modules of Aurora and Polo-Like Kinase Activities in Mitotic Cells. Science Signaling, 179:rs5.

Krenn, V., and A. Musacchio. 2015. The Aurora B kinase in chromosome bi-orientation and spindle checkpoint signaling. Front Oncol. 5:225.

Lampert, F., Hornung, P., and Westermann, S. 2010. The Dam1 complex confers microtubule plus end-tracking activity to the Ndc80 kinetochore complex. Journal of Cell Biology, 189:641–649.

Lampert, F., Mieck, C., Alushin, G. M., Nogales, E., and Westermann, S. 2013. Molecular requirements for the formation of a kinetochore-microtubule interface by Dam1 and Ndc80 complexes. Journal of Cell Biology, 200:21–30.

Manning, A. L., and Compton, D. A. 2007. Mechanisms of Spindle-Pole Organization Are Influenced by Kinetochore Activity in Mammalian Cells. Current Biology, 17:260–255.

Maiato, H., and Logarinho, E. 2014. Mitotic spindle multipolarity without centrosome amplification. Nature Cell Biology, 16:386–394.

Maiolica, A., Cittaro, D., Borsotti, D., Sennels, L., Ciferri, C., Tarricone, C., Musacchio, A., and Rappsilber, J. 2007. Structural Analysis of Multiprotein Complexes by Cross-linking, Mass Spectrometry, and Database Searching. Molecular & Cellular Proteomics, 6.12: 2200–2211

Mattiuzzo, M., Vargiu, G., Totta, P., Fiore, M., Ciferri, C., Musacchio, A., and Degrassi, F. 2011. Abnormal Kinetochore-Generated Pulling Forces from Expressing a N-Terminally Modified Hec1. PLoS ONE, 6:e16307.

Miller, S. A., Johnson, M. L., and Stukenberg, P. T. 2008. Kinetochore Attachments Require an Interaction between Unstructured Tails on Microtubules and Ndc80^Hec1^. Curr Biol, 18:1785–1791.

Nousiainen, M., Sillje, H., Sauer, G., Nigg, E. A., and Ko, R. 2006. Phosphoproteome analysis of the human mitotic spindle. PNAS, 103:5391–5396.

Powers, A. F., Franck, A. D., Gestaut, D. R., Cooper, J., Gracyzk, B., Wei, R. R., Wordeman, L., Davis, T. N., and Asbury, C. L. 2009. The Ndc80 Kinetochore Complex Forms Load-Bearing Attachments to Dynamic Microtubule Tips via Biased Diffusion. Cell, 136:865–875.

Raaijmakers, J. A., Tanenbaum, M. E., Maia, A. F., and Medema, R. H. 2009. RAMA1 is a novel kinetochore protein involved in kinetochore-microtubule attachment. Journal of Cell Science, 122:2436–2445.

Roscioli E., Germanova, T. E., Smith, C. A., Embacher, P. A., Erent, M., Thompson, A. I., Burroughs, N. J., and McAinsh, A. D. 2019. Ensemble-level organization of human kinetochores and evidence for distinct tension and attachment sensors. BioRxiv.

Schmidt, J. C., Arthanari, H., Boeszoermenyi, A., Natalia, M., Wilson-Kubalek, E. M., Monnier, N., Markus, M., Oberer, M., Milligan, R. A., Bathe, M., Wagner, G., Grishchuk, E. L., and Cheeseman, I. M. 2013. The kinetochore-bound Ska1 complex tracks depolymerizing microtubules and binds to curved protofilaments. Developmental Cell, 23:968–980.

Sivakumar, S., Janczyk, P. Ł., Qu, Q., Brautigam, C. A., Stukenberg, P. T., Yu, H., and Gorbsky, G. J. 2016. The human SKA complex drives the metaphase-anaphase cell cycle transition by recruiting protein phosphatase 1 to kinetochores. eLIFE. 5:e12902.

Sivakumar, S., Daum, J.R., Tipton, A. R., Rankin, S., and Gorbsky, G.J. 2014. The spindle and kinetochore-associated (Ska) complex enhances binding of the anaphase-promoting complex/cyclosome (APC/C) to chromosomes and promotes mitotic exit. Mol Biol Cell. 25:594–605.

Sundin, L. J. R., Guimaraes, G. J., and DeLuca, J. G. 2011. The NDC80 complex proteins Nuf2 and Hec1 make distinct contributions to kinetochore-microtubule attachment in mitosis. Molecular Biology of the Cell, 22:759–768.

Suzuki, A., Badger, B. L., Haase, J., Ohashi, T., Erickson, H. P., Salmon, E. D., and Bloom, K. 2016. How the kinetochore couples microtubule force and centromere stretch to move chromosomes. Nature Cell Biology, 18:382–392.

Suzuki, A., Long, S. K., and Salmon, E. D. 2018. An optimized method for 3D fluorescence co-localization applied to human kinetochore protein architecture. eLife, 7:e32418.

Tanaka, T.U., N. Rachidi, C. Janke, G. Pereira, M. Galova, E. Schiebel, MJR. Stark, and K. Nasmyth. 2002. Evidence that the Ipl1-Sli5 (Aurora kinase-INCENP) complex promotes chromosome bi-orientation by altering kinetochore-spindle pole connections. Cell. 108: 317–329.

Tauchman, E.C., Boehm, F. J., and DeLuca, J. G. 2015. Stable kinetochore-microtubule attachment is sufficient to silence the spindle assembly checkpoint in human cells. Nat Commun. 6:10036.

Theis, M., Slabicki, M., Junqueira, M., Paszkowski, M., Sontheimer, J., Kittler, R., Heninger, A., Glatter, T., Kruusmaa, K., Poser, I., Hyman, A. A., Pisabarro, M. T., Gstaiger, M., Aebersold, R., Shevchenko, A., and Buchholz, F. 2009. Comparative profiling identifies C13orf as a component of the Ska complex required for mammalian cell division. The EMBO Journal, 28:1453–1465.

Tien, J. F., Umbreit, N. T., Gestaut, D. R., Franck, A. D., Cooper, J., Wordeman, L., Gonen, T., Asbury, C. L., and Davis, T. N. 2010. Cooperation of the Dam1 and Ndc80 kinetochore complexes enhances microtubule coupling and is regulated by aurora B. Journal of Cell Biology, 189:713–723.

Tooley, J. G., Miller, S. A., Stukenberg, P. T., and Drubin, D. G. 2011. The Ndc80 complex uses a tripartite attachment point to couple microtubule depolymerization to chromosome movement. Molecular Biology of the Cell, 22:1217–1226.

Umbreit, N. T., Gestaut, D. R., Tien, J. F., Vollmar, B. S., Gonen, T., and Asbury, C. L. 2012. The Ndc80 kinetochore complex directly modulates microtubule dynamics. PNAS, 109:16113–16118.

Varma, D., Chandrasekaran, S., Sundin, L. J. R., Reidy, K. T., Wan, X., Chasse, D. A. D., Nevis, K. R., DeLuca, J. G., Salmon, E. D., and Cook, J. G. 2012. Recruitment of the human Cdt1 replication licensing protein by the loop domain of Hec1 is required for stable kinetochore-microtubule attachment. Nature Cell Biology, 14:593–603.

Varma, D., and Salmon, E. D. 2012. The KMN protein network—chief conductors of the kinetochore orchestra. Journal of Cell Science, 125:5927–5936.

Wan, X., O’Quinn, R. P., Pierce, H. L., Joglekar, A. P., Gall, W. E., DeLuca, J. G., Carroll, C. W., Liu, S. T., Yen, T. J., McEwen, B. F., Stukenberg, P. T., Desai, A., and Salmon, E. D. 2009. Protein architecture of the human kinetochore microtubule attachment site. Cell. 137:672–684.

Wei, R. R., Al-Bassam, J., and Harrison, S. C. 2007. The Ndc80/HEC1 complex is a contact point for kinetochore-microtubule attachment. Nature Structural & Molecular Biology, 14:54–59.

Welburn, J. P. I., Grishchuk, E. L., Backer, C. B., Wilson-Kubalek, E. M., Yates III, J. R., and Cheeseman, I. M. 2009. The Human Kinetochore Ska1 Complex Facilitates Microtubule Depolymerization-Coupled Motility. Developmental Cell, 16:374–385.

Wilson-Kubalek, E. M., Cheeseman, I. M., Yoshioka, C., Desai, A., and Milligan, R. A. 2008. Orientation and structure of the Ndc80 complex on the microtubule lattice. Journal of Cell Biology, 182:1055–1061.

Yoo, T.Y., J. Choi, W. Conway, C. Yu, RV. Pappu, and DJ. Needleman. 2018. Measuring NDC80 binding reveals the molecular basis of tension-dependent kinetochore-microtubule attachments. Elife. 7:1–34.

Zaytsev, AV., LJR. Sundin, KF. DeLuca, EL. Grishchuk, JG. and DeLuca. 2014. Accurate phosphoregulation of kinetochore-microtubule affinity requires unconstrained molecular interactions. J Cell Biol. 206:45–59.

Zaytsev, A. B., Mick, J. E., Maslennikov, E., Nikashin, B., DeLuca, J. G., and Grishchuk, E. L. 2015. Multisite phosphorylation of the NDC80 complex gradually tunes its microtubule-binding affinity. Molecular Biology of the Cell, 26:1829–1844.

Zhai, Y., Kronebusch, P. J., and Borisy, G. G. 1995. Kinetochore Microtubule Dynamics and the Metaphase-Anaphase Transition. Journal of Cell Biology, 131:721–734.

Zhang, G., Kelstrup, C. D., Hu, X., Hansen, M. J. K., Singleton, M. R., Olsen, J. V., and Nilsson, J. 2012. The Ndc80 internal loop is required for recruitment of the Ska complex to establish end-on microtubule attachment to kinetochores. Journal of Cell Science, 135:3243–3253.

Zhang, Q., Chen, Y., Yang, L., and Liu, H. 2018. Multitasking Ska in Chromosome Segregation: Its Distinct Pools Might Specify Various Functions. BioEssays 40:1700176.

Zhang, Q., Sivakumar, S., Chen, Y., Gao, H., Yang, L., Yuan, Z., Yu, H., and Liu, H. 2017. Ska3 Phosphorylated by Cdk1 Binds Ndc80 and Recruits Ska to Kinetochores to Promote Mitotic Progression. Curr Biol, 27:1477–1484.

