## Supplemental Material for "Coordination of NDC80 and Ska complexes at the kinetochore-microtubule interface in human cells"

**Supplemental Figure Legends**

**Supplemental Figure 1. Expression of 9A-Hec1-GFP in cells depleted of endogenous Hec1 increases Ska complex loading to kinetochores.** (A) Immunofluorescence images of WT- and 9A-expressing cells depleted of endogenous Hec1 with siRNA and stained with antibodies to Hec1 pS69 and Ska3. (B) Quantification of pS69 kinetochore fluorescence intensity from Hec1 siRNA-treated cells expressing WT- and 9A-Hec1-GFP. For each condition, at least 20 kinetochores per cell were measured from at least 9 cells per experiment from 2 separate experiments. (C) Quantification of Ska3 kinetochore fluorescence intensity from Hec1 siRNA-treated cells expressing WT- and 9A-Hec1-GFP. For each condition, at least 20 kinetochores per cell were measured from at least 9 cells per experiment from 2 separate experiments. A Student's t-test was carried out to determine statistical significance.

**Supplemental Figure 2. Asynchronous cells expressing non-phosphorylatable Hec1 exhibit increased levels of Ska3 after nocodazole treatment** (A) Immunofluorescence images of asynchronous cells expressing WT- and 9A-Hec1-GFP treated with or without 10 µm nocodazole for 1h prior to fixation. Cells were fixed and stained with antibodies to Ska3 (rabbit). (B) Quantification of Ska3 kinetochore fluorescence intensity from cells expressing WT- and 9A-Hec1-GFP treated with or without nocodazole. For each condition, at least 20 kinetochores per cell were measured from at least 5 cells per experiment from 4 separate experiments. Statistical significance was determined by a one-way Anova analysis. (C) Immunofluorescence panels of control, untreated HeLa cells or HeLa cells treated with Ska1 and Ska3 siRNA. Cells were incubated in cold DMEM on ice for 12 minutes prior to fixation, permeabilized, fixed, and stained with antibodies to tubulin and an anti-centromere-antibody (ACA). Insets are enlargements of the region indicated by the dashed box. (D) Quantification of end-on attachment in control cells and cells treated with Ska1 and Ska3 siRNA. For each condition, at least 15 kinetochores per cell were measured from 10 cells per experiment from 2 separate experiments.

**Supplemental Figure 3. Oligomerization of Ska and NDC80 complexes *in vitro* is buffer dependent.** (A) GFP fluorescence (top row) and overlay with Alexa647-tubulin (bottom row) images of GFP-tagged Ska complex (SkaC-GFP) diluted to the noted concentrations in buffers indicated above each column. SkaC-GFP microtubule binding reactions were carried out in the same manner as experiments from Figure 3 (see Materials and Methods for detailed information). (B) GFP fluorescence (top row) and overlay with Alexa647-tubulin (bottom row) images of indicated NDC80C-GFP constructs incubated with unlabeled Ska complex (SkaC) in BRB80 buffer.

**Supplemental Figure 4. Hec1 tail deletion impacts tension generation at kinetochores.** (A) Immunofluorescence images of HeLa cells expressing WT- and Δ80-Hec1-GFP. Cells were incubated in cold DMEM on ice for 12 minutes prior to fixation, permeabilized, fixed, and stained with antibodies to tubulin. Insets are enlargements of the region indicated by the dashed box. (B) Quantification of inter-kinetochore distances in metaphase and prometaphase cells expressing WT-Hec1-GFP, and cells expressing Δ80-Hec1-GFP. For each condition, inter-kinetochore distances were measured from at least 110 kinetochores per cell in at least 11 cells per experiment from at least 3 independent experiments. (C) Immunofluorescence images of HeLa cells expressing WT- and Δ80-Hec1-GFP and depleted of endogenous Hec1 stained with antibodies to tubulin. Insets are enlargements of the region indicated by the dashed box. (D) Quantification of chromosome alignment in cells expressing WT- and Δ80-Hec1-GFP. For each condition, chromosome alignment was assessed in at least 100 cells per experiment in 2 separate experiments. Cells were scored as "aligned" if they had a metaphase plate with <5 chromosomes off the plate. (E) Quantification of multipolarity observed in cells expressing WT- and Δ80-Hec1-GFP. Cells with unaligned chromosomes were scored for containing bi- vs multi-polar spindles, and the percent of cells with multipolar spindles is shown. For each condition, at least 100 cells per experiment from two separate experiments were assessed.

**Supplemental Figure 5. Location of the GFP tag differentially affects ability of cells expressing Δ80-Hec1 to form stable kinetochore-microtubule attachments.** (A) Immunofluorescence images of HeLa cells expressing N- and C-terminally GFP-tagged WT- and Δ80-Hec1 constructs. Insets are enlargements of the region indicated by the dashed box. Schematics of the constructs used are indicated on the right. (B) Immunofluorescence images of cold-treated HeLa cells expressing N- and C-terminally tagged Δ80-Hec1 constructs. Cells were incubated in cold DMEM on ice for 12 minutes prior to fixation, permeabilized, fixed, and stained with antibodies to tubulin. Insets are enlargements of the region indicated by the dashed box. (C) Quantification of end-on kinetochore-microtubule attachment in cold-treated HeLa cells expressing WT-Hec1-GFP, and N- and C-terminally tagged Δ80-Hec1 constructs. For N- and C-terminally tagged Δ80-Hec1 constructs, at least 15 kinetochores per cell were quantified from at least 8 cells from 3 independent experiments. For the WT-Hec1-GFP metaphase data, 15 kinetochores per cell were quantified from 5 cells per experiment from 2 independent experiments. The C-terminally tagged Δ80-Hec1WT-Hec1-GFP data shown are from the experiment presented in Figure 4. Statistical significance was determined using a Student's t-test. (D) Immunofluorescence images of cold-treated RPE1 cells expressing N- and C-terminally GFP-tagged Δ80-Hec1 constructs. Cells were incubated in cold DMEM on ice for 15 minutes prior to fixation, permeabilized, fixed, and stained with antibodies to tubulin. Insets are enlargements of the region indicated by the dashed box. Schematics of the constructs used are indicated on the right.

**Supplemental Figure 6. Specific regions of the Hec1 loop are required for chromosome alignment in cells.** (A) List of loop mutants used. (B) Immunofluorescence images of cells expressing WT- or ML-Hec1-GFP constructs and stained with Astrin and tubulin antibodies. (C) Quantification of chromosome alignment in cells expressing WT- or ML-Hec1-GFP constructs. For each condition, at least 120 cells were analyzed from at least 2 separate experiments. Cells were scored as “aligned” if they had a metaphase plate with <5 chromosomes off the plate.

Wimbish et al., Figure S1

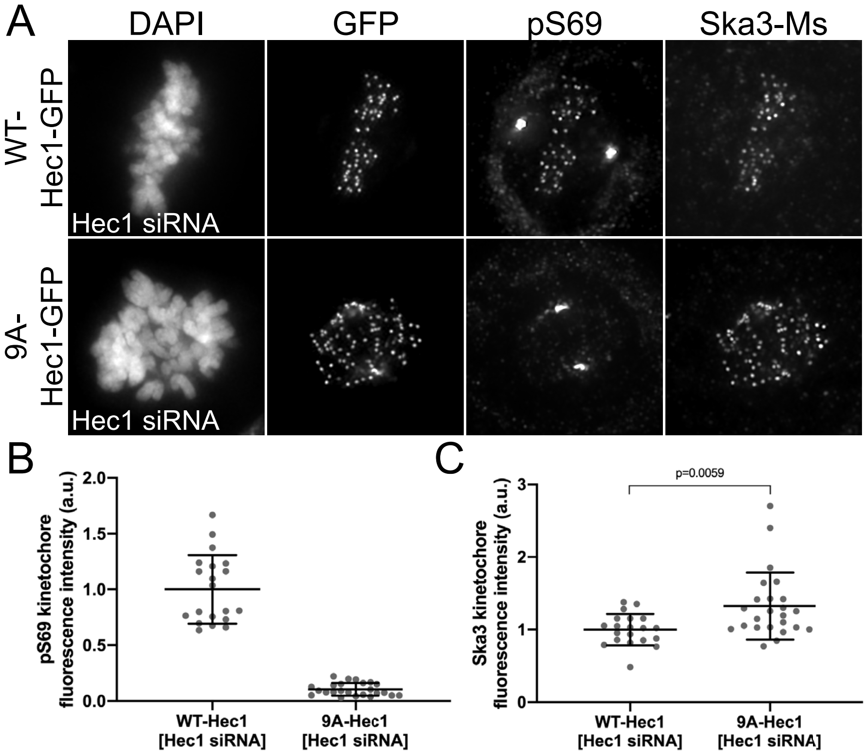

Wimbish et al., Figure S2

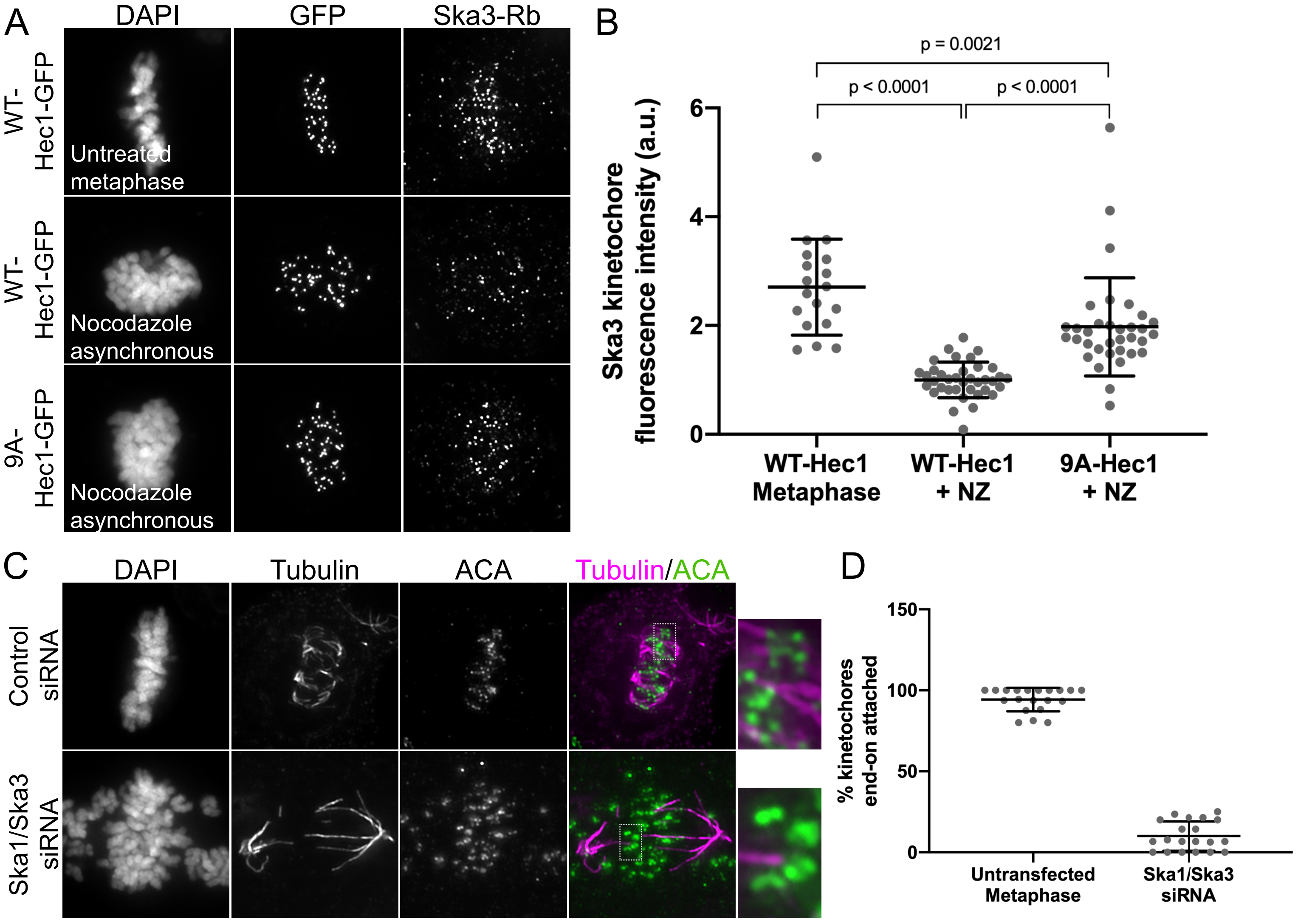

Wimbish et al., Figure S3

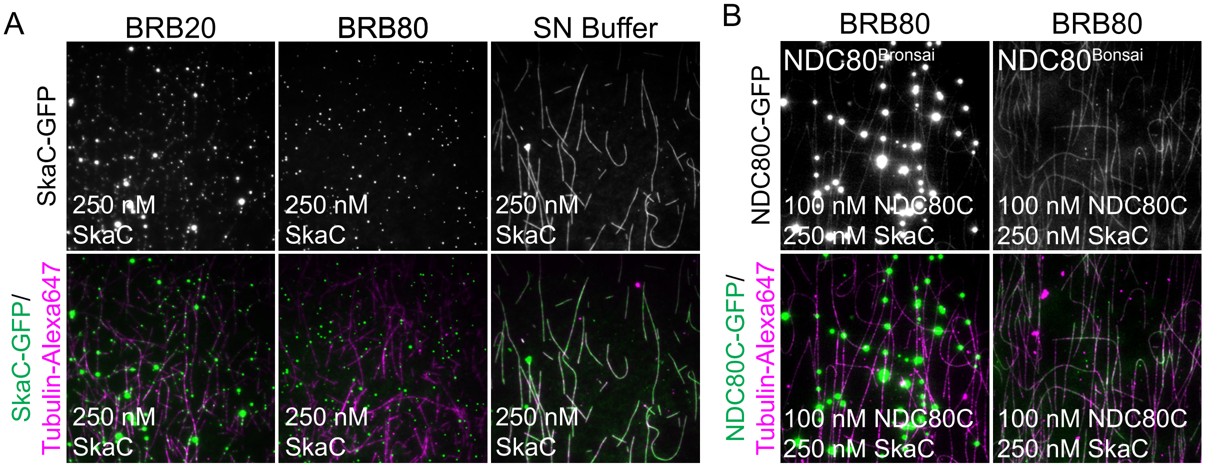

### Wimbish et al., Figure S4

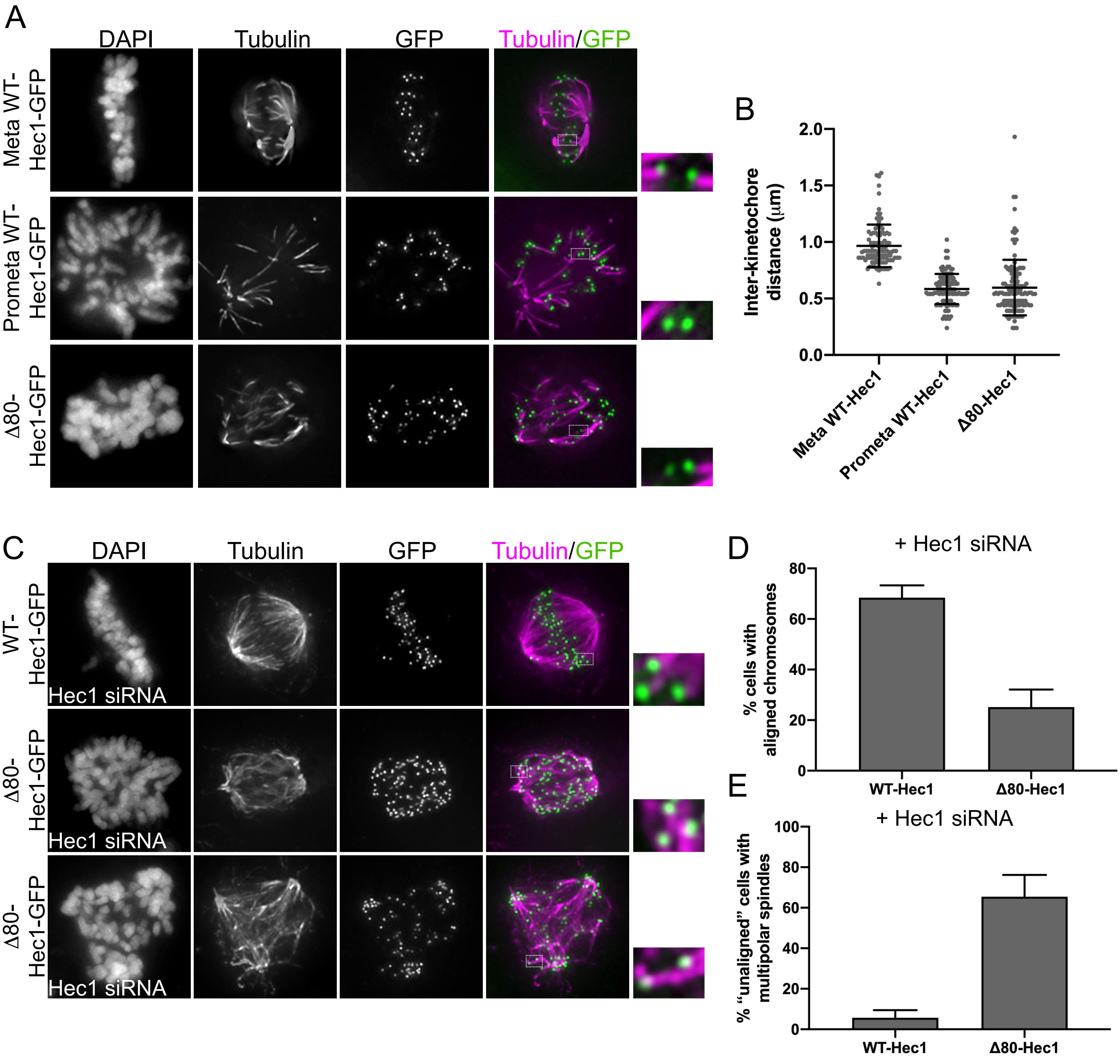

Wimbish et al., Figure S5

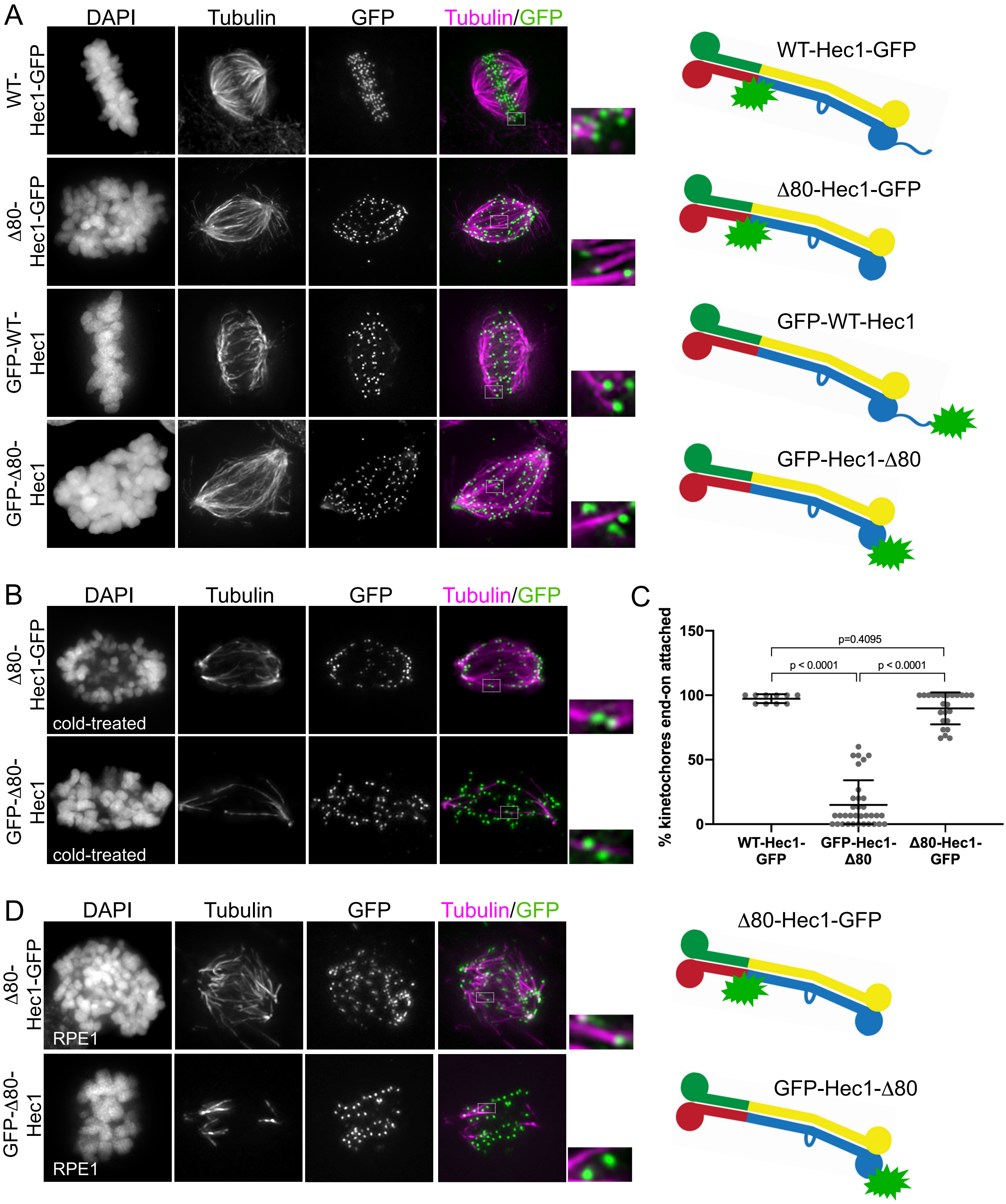

Wimbish et al., Figure S6

A

WT : ***420*** RKLKLIPKGAENSKGYDFEIKFNPEAGANCLVKYRAQVYVP ***460***

ML$^{420-424A}$: ***420*** **AAAAA**IPKGAENSKGYDFEIKFNPEAGANCLVKYRAQVYVP ***460***

ML$^{425-429A}$: ***420*** RKLKL**AAAAA**ENSKGYDFEIKFNPEAGANCLVKYRAQVYVP ***460***

ML$^{430-434A}$: ***420*** RKLKLIPKGA**AAAAA**YDFEIKFNPEAGANCLVKYRAQVYVP ***460***

ML$^{435-439A}$: ***420*** RKLKLIPKGAENSKG**AAAAA**KFNPEAGANCLVKYRAQVYVP ***460***

ML$^{440-444A}$: ***420*** RKLKLIPKGAENSKGYDFEI**AAAAA**AGANCLVKYRAQVYVP ***460***

ML$^{445-449A}$: ***420*** RKLKLIPKGAENSKGYDFEIKFNPE**AAAAA**LVKYRAQVYVP ***460***

ML$^{450-454A}$: ***420*** RKLKLIPKGAENSKGYDFEIKFNPEAGANC**AAAAA**AQVYVP ***460***

ML$^{456-460A}$: ***420*** RKLKLIPKGAENSKGYDFEIKFNPEAGANCLVKYR**AAAAAA** ***460***

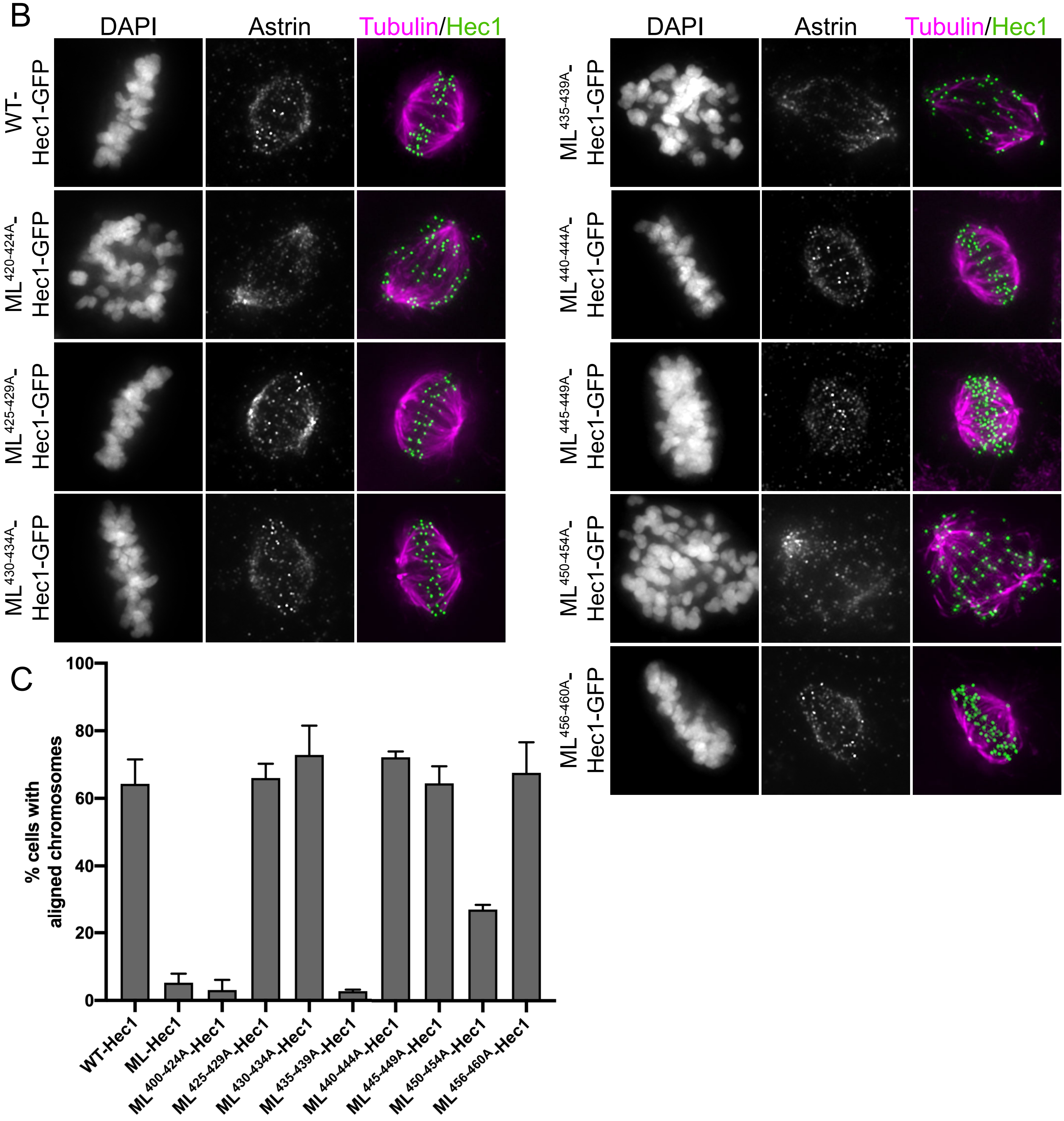
